# Selective Vulnerability of Layer 5a Corticostriatal Neurons in Huntington’s Disease

**DOI:** 10.1101/2023.04.24.538096

**Authors:** Christina Pressl, Kert Mätlik, Laura Kus, Paul Darnell, Ji-Dung Luo, Matthew R. Paul, Alison R. Weiss, William Liguore, Thomas S. Carroll, David A. Davis, Jodi McBride, Nathaniel Heintz

## Abstract

The properties of the cell types that are selectively vulnerable in Huntington’s disease (HD) cortex, the nature of somatic CAG expansions of *mHTT* in these cells, and their importance in CNS circuitry have not been delineated. Here we employed serial fluorescence activated nuclear sorting (sFANS), deep molecular profiling, and single nucleus RNA sequencing (snRNAseq) to demonstrate that layer 5a pyramidal neurons are vulnerable in primary motor cortex and other cortical areas of HD donors. Extensive *mHTT*-CAG expansions occur in vulnerable layer 5a pyramidal cells, and in Betz cells, layer 6a, layer 6b neurons that are resilient in HD. Retrograde tracing experiments in macaque brains identify the vulnerable layer 5a neurons as corticostriatal pyramidal cells. We propose that enhanced somatic *mHTT*-CAG expansion and altered synaptic function act together to cause corticostriatal disconnection and selective neuronal vulnerability in the HD cerebral cortex.

## INTRODUCTION

Neurodegeneration in Huntington’s disease (HD) is evident primarily in the striatum in early stages, and it is thought that loss of medium–sized spiny neurons (MSNs) in the caudate nucleus and putamen is responsible for motor abnormalities that are cardinal features of the disease (Reiner and Deng, 2018; Vonsattel et al., 2011; Vonsattel et al., 1985). The clinical appearance of motor symptoms is used as a measure of age at onset, which is inversely correlated with the number of CAG repeats in exon 1 of the mutant *HTT* allele (*mHTT*), often defined by the CAG tract length in blood (Andrew et al., 1993; Hong et al., 2021; Monckton, 2021; Ross and Tabrizi, 2011). Neuroimaging studies of the human cerebral cortex have demonstrated that connectivity between cortical areas and the basal ganglia is disturbed very early in the HD individuals (Espinoza et al., 2018; Gargouri et al., 2016; McColgan et al., 2017) (Jernigan et al., 1991) (Hasselbalch et al., 1992; Jenkins et al., 1998; Kuhl et al., 1982; Kuwert et al., 1989, 1990; Sax et al., 1996; Unschuld et al., 2012), and that widespread cortical degeneration occurs as the disease progresses (Johnson et al., 2021; Minkova et al., 2018; Odish et al., 2018; Tabrizi et al., 2013). Although *HTT* is ubiquitously expressed (Landwehrmeyer et al., 1995), substantial variations in mood, motor function, and cognition in HD are associated with region specific neuropathological findings occurring in the prefrontal, motor and cingulate cortices (Hickman et al., 2022; Mehrabi et al., 2016; Nana et al., 2014; Nopoulus, 2010). Moreover, results from functional connectivity imaging studies have suggested that alterations in the connectivity within the indirect and direct basal ganglia pathways differentially affect symptoms in HD patients (Nair et al., 2022). Genome-wide association studies indicate that the effect of HD modifying genes influence the motor and cognitive domains of the disease differentially (Correia et al., 2015; Lee et al., 2022; Moss et al., 2017). Evidence of altered cortical function has also been documented in a variety of HD mouse models (Gu et al., 2022; MacDonald et al., 2003; Veldman and Yang, 2018) including changes in electrophysiological properties of cortical projection neurons (Blumenstock and Dudanova, 2020; Cepeda et al., 2010) although important differences in cortical function have been noted between these models.

Despite the wealth of information on cortical dysfunction and degeneration that has been obtained in studies of HD, our knowledge of histological and molecular events that occur in early stages of the disease is incomplete. It is not known, for example, whether the somatic expansion of the *mHTT*- exon1 CAG tract that has been demonstrated in the cortex (Shelbourne et al., 2007; Swami et al., 2009) occurs specifically in vulnerable neurons, or how the molecular pathology of each cortical cell type differs as the disease progresses.

To address these issues, we have developed serial fluorescence activated nuclear sorting (sFANS) of cortical samples to enable deep molecular analysis of cortical neuron classes from control and HD donors. Our focused studies of the primary motor cortex (BA4) using sFANS, supplemented with single nucleus RNA seq (snRNAseq), demonstrate that Layer 5a (L5a) pyramidal cells are selectively vulnerable in the human motor cortex in early HD. Analysis of deep layer pyramidal neuron samples from a total of six control donors and thirteen HD donors establish also that loss of Layer 5a neurons occurs widely in the cerebral cortex in HD donors. Although expansion of the *mHTT* exon1 CAG tract to ∼60-100 repeats is evident in vulnerable layer 5a cells in the HD cortex, equally robust somatic CAG expansions are also present in Betz cells, Layer 6a and Layer 6b pyramidal neurons that remain resilient during the early phases of HD degeneration. Retrograde tracing using adeno associated viral (AAV) vectors to identify striatal projecting neurons in macaque brain (Weiss et al., 2020), coupled with *in situ* hybridization probes specific for the selectively vulnerable layer 5a human cortical neurons, strongly support the conclusion that corticostriatal projection neurons are especially vulnerable in HD. These data are consistent with identification of transcriptional programs preferentially impacted in Layer 5a neurons that indicate synaptic functions may be altered in corticostriatal projection neurons in HD. Taken together, our data identify striatal projecting Layer 5a neurons as vulnerable in multiple cortical regions in HD, they demonstrate that extensive somatic CAG expansions occur in both vulnerable and more resilient deep layer cortical neurons, and they suggest that synaptic dysfunction and the cortico-striatal disconnection documented in prior human studies (McColgan et al., 2017) may play an important role in HD neurodegeneration.

## RESULTS

### sFANSseq for profiling the major cell classes of the human cerebral cortex

To begin to understand the molecular pathology that is associated with Huntington’s disease in the human cerebral cortex, we sought to develop an approach that could provide comprehensive transcriptional data filtered for promoter accessibility as an independent measure of active genes, as well as somatic CAG tract expansion measurements of the *mHTT* allele, from isolated populations of nuclei representing the major cell classes. Given the requirement for very efficient use of small tissue samples, especially given the limited availability of samples from HD donors, we built on the fluorescence activated nuclear sorting (FANS) strategy (Xu et al., 2018). Identification of a cortex-specific multiplexed antibody panel and the discovery that a serial sorting and serial staining strategy was optimal for high resolution characterization of all major cortical cell types from a single sample in a single session allowed us to achieve these aims. Using this serial fluorescence activated nuclear sorting (sFANS) strategy (Figure 1A), we were able to isolate nuclei from 14-16 distinct cell classes depending on the cortical region being analyzed.

**Figure 1.**
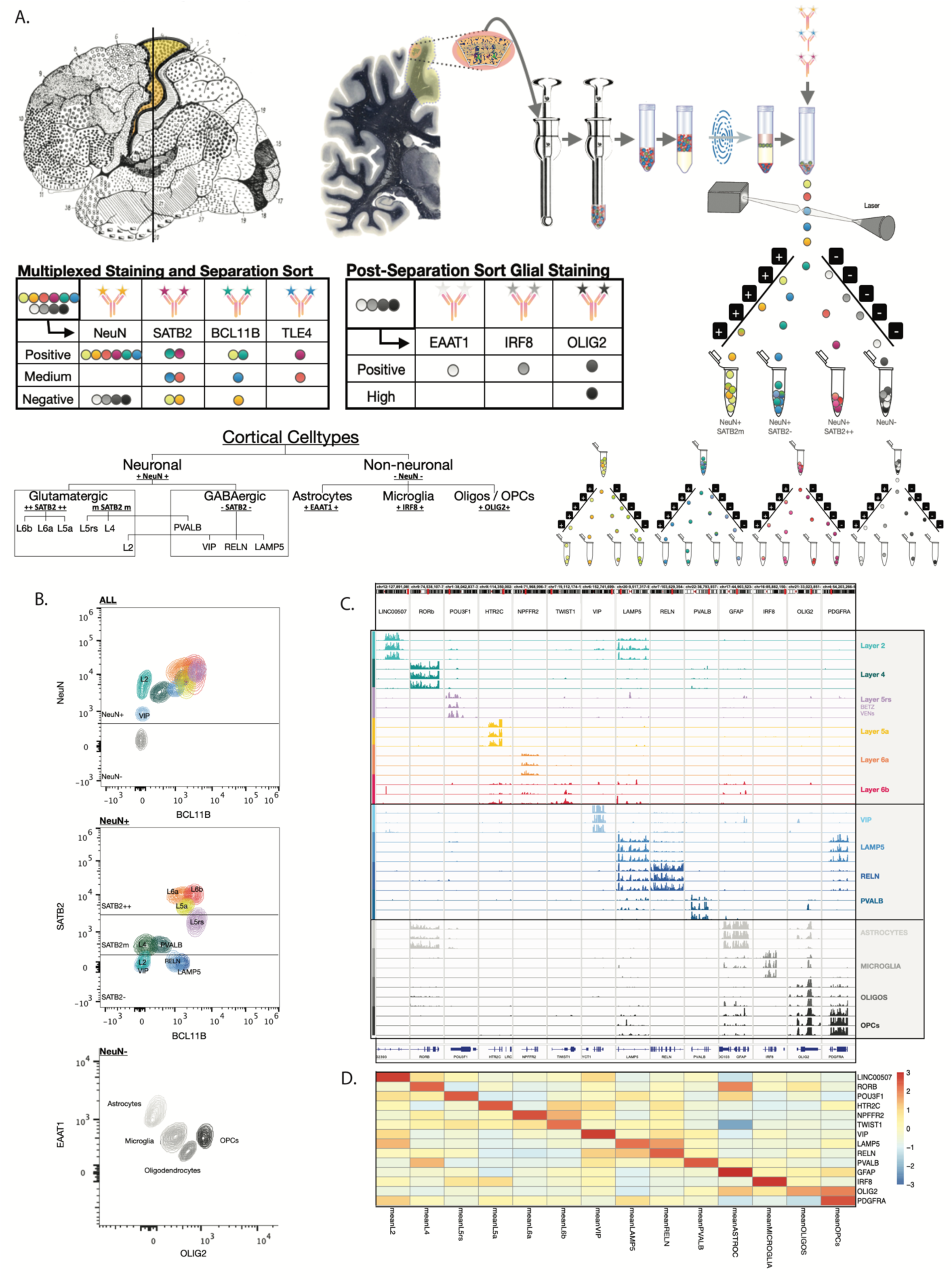
sFANSseq for profiling the major cell classes of the human cerebral cortex. (**A**) Schematic representation of human cortical sample preparation, nuclei extraction, and serial fluorescence activated nuclei sorting (sFANS), using multiplexed fluorescent antibody panels. (**B**) Representative sFANS plots from one motor cortex sample, showing distribution of the major human cortical cell-type populations. (**C**) Examples of resulting sFANSseq IGV gene expression profiles from multiple regions of cortex and donors. Populations shown include layer 2 (L2), layer 4 (L4), layer 5 region-specific neurons (L5rs), layer 5a (L5a), layer 6a (L6a), layer 6b (L6b), VIP expressing interneurons, LAMP5 expressing interneurons (LAMP5), RELN expressing interneurons (RELN), PVALB expressing interneurons (PVALB), astrocytes, microglia, oligodendrocytes (Oligos), and oligodendrocyte progenitor cells (OPCs). (**D**) Heatmap shows log-2 transformed average DESeq-normalized expression levels for cell-type specific marker genes across all samples.

To identify the cognate cell class for each sFANS sorted population, RNA-sequencing was employed to generate nuclear gene expression profiles through sFANSseq. The sFANS sorting parameters and analysis of marker gene expression in the isolated nuclei identified the recovered sFANS nuclei as: excitatory neurons from Layer 2 (L2), Layer 4 (L4), Layer 5rs (region-specific; L5rs), Layer 5a (L5a), Layer 6a (L6a), Layer 6b (L6b); inhibitory interneurons expressing VIP, LAMP5, RELN, and PVALB; as well as the non-neuronal cells astrocytes, microglia, oligodendrocytes, and oligodendrocyte progenitor cells (Figure 1B). The sFANS strategy was effective for analysis of these cell types in six regions of the human cortex, including the motor cortex (BA4), prefrontal cortex (BA9), visual cortex (BA17-19), auditory cortex (BA22), insular cortex (INS, BA13-16), and cingulate cortex (BA24). Furthermore, variations in the sFANS profiles in samples from specific cortical areas allowed identification and characterization of rare and region-specific cell types, including Betz cells (Bakken et al., 2021), Von Economo neurons (Hodge et al., 2020), and CRHBP expressing interneurons (Li et al., 2016) (Supplementary Figure S1A-C). Successful enrichment for cell type specific markers in the expression profiles (Figure 1C), as well as the gene expression heatmap of these data (Figure 1D) and principal component analyses (Supplementary Figure S1D) demonstrate the reproducibility and resolution of the sFANS approach. Although further resolution of some of these cell classes into subclasses evident in snRNAseq data (Bakken et al., 2021) can be achieved using subclass specific probes for targeted sFANS isolation, the sFANS approach was used routinely in this study because it can provide comprehensive molecular data on all cortical cell classes in a single session for each cortical sample. Using the sFANS approach we used 42 NC samples for our specificity index analyses, for our deep layer focused RNAseq differential gene expression analyses we utilized 85 NC and 78 HD samples. In our ATACseq analyses 16 NC samples from four deep layer cell types were used. Nine NC and nine HD samples were used for the targeted snRNAseq analyses and lastly, 148 HD samples were utilized for CAG repeat expansion analyses (for more details see Table S2).

### Layer 5a 5-Hydroxytryptamine Receptor 2C (*HTR2C*) expressing pyramidal cells are selectively vulnerable in HD motor cortex

To study the impact of HD on each of the cell classes identified in sFANS, we used this strategy to characterize nuclei isolated from control and HD donor samples. Analysis of the sFANS data from control samples revealed clear separation of 3 distinct subpopulations expressing high levels of the excitatory neuron markers Special AT-rich sequence-binding protein 2 (*SATB2*) (Alcamo et al., 2008) and Transducin-like enhancer protein 4 (*TLE4*) (Hevner, 2007) (Figure 2A). Analysis of transcriptional profiles collected from each of these populations relative to snRNAseq data (Bakken et al., 2021) established that a healthy *SATB2* high population contains three subpopulations of deep layer glutamatergic cell-types: *HTR2C* expressing Layer 5a neurons, as well as two separate Cellular Communication Network Factor 2 (*CCN2*) expressing deep layer 6 populations expressing Neuropeptide FF Receptor 2 (*NPFFR2*) and Sulfatase 1 (*SULF1*), respectively. Recovery and analysis of Betz cells from the motor cortex was evident also from marker expression, revealing a distinct population of large nuclei expressing POU Domain Transcription Factor (*POU3F1*) (Bakken et al., 2021) in the *SATB2* medium range of the sFANS data (Supplementary Figure S1A and Supplementary Figure S2B).

**Figure 2.**
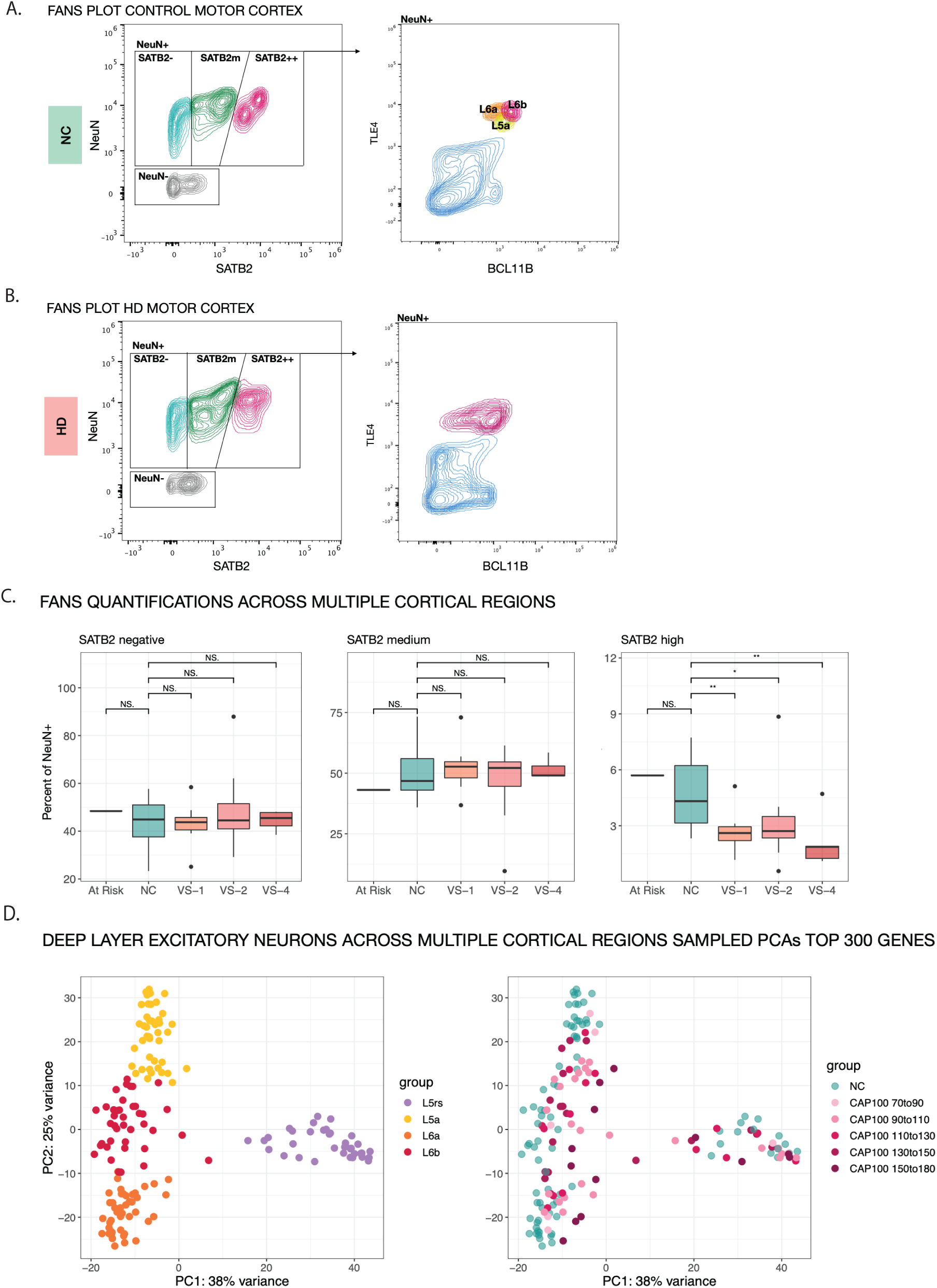
Layer 5a *HTR2C* expressing pyramidal cells are selectively vulnerable in HD motor cortex. (**A**) Representative sFANS panels from one control donor showing distribution of NeuNnegative, NeuNpositive/SATB2negative, NeuNpositive/SATB2medium, and NeuNpositive/SATB2high populations (left panel). Representative distribution of NeuNpositive nuclei plotted for fluorophore signal intensity of TLE4 and BCL11b, revealing three TLE4 positive deep layer excitatory neuron populations layer 5a (L5a), layer 6a (L6a), layer 6b (L6b) (right panel). (**B**) Representative sFANS panels from one HD donor showing distribution of NeuNnegative, NeuNpositive/SATB2negative, NeuNpositive/SATB2medium, and NeuNpositive/SATB2high populations (left panel). Representative distribution of NeuNpositive nuclei plotted for fluorophore signal intensity of TLE4 and BCL11b, revealing a distortion of the staining pattern of TLE4 positive populations (right panel). (**C**) Quantitation of the relative abundance of NeuNpositive/SATB2negative, NeuNpositive/SATB2medium, and NeuNpositive/SATB2high nuclei among all NeuNpositive nuclei isolated from three Normal Control (NC) donors, one genetically confirmed HD donor pre-diagnosis (At Risk), four HD donors with Vonsattel striatal grade 1 (VS-1), three HD donors with Vonsattel striatal grade 2 (VS-2), one HD donor with Vonsattel striatal grade 4 (VS-4). (**D**) Principal component analyses (PCA) utilizing a total of 163 datasets from the four deep layer excitatory neuron populations layer 5 region-specific neurons (L5rs), layer 5a (L5a), layer 6a (L6a), layer 6b (L6b), color coded by cell-type (left panel) and color coded by group (right panel), with controls in green and HD samples in shades of red, grouped by the normalized CAG-age product (CAP100) (CAP100 70to90, CAP100 90to110, CAP100 110to130, CAP100 130to150, CAP100 150to180). CAP100 scores were calculated as described by Warner et al. (Warner et al., 2022):

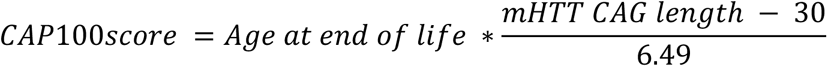

Analysis of sFANS plots from HD donor samples revealed a clear distortion in the staining pattern of nuclei expressing high levels of *SATB2* and *TLE4* (Figure2B, Supplementary Figure S2A). To quantify this observation, the relative number of nuclei stained in the initial sFANS as NeuN positive and SATB2 negative, medium, and high fluorescence was plotted for three control vs three HD donors (Figure 2C). These data revealed a decrease of SATB2 high nuclei from HD donors with a range of Vonsattel grades (VS) (Vonsattel et al., 1985), a first indication of HD-specific cell loss in the deep layer populations. To understand which deep layer pyramidal cell class was most impacted in the HD samples, we used PCA of sFANSseq datasets from the four deep layer neuronal populations. Labeling the PCA plots according to cell type revealed each of the deep layer neuronal types in discrete areas of the plot (Figure 2D, left). However, labeling by donor identity and CAP score revealed that very few samples from HD brains clustered in the layer 5a domain of the PCA plot evident from control donors (Figure 2D, right). This finding was evident in 163 cell-type specific sFANSseq samples, including 85 NC and 78 HD cell-type specific samples, derived from both male and female donors, including 6 NC and 13 HD donors (Number of male NC = 65, female NC = 20, male HD = 38, female HD = 40 donor samples. The number of cell-type specific samples across HD donor cap scores ranges were: CAP100 70-90=7, CAP100 90-110=25; CAP100 110-130=15; CAP100 130-150=18; and CAP100 150-180=13 (for more details see Table S2). It is apparent from these data that recovery of layer 5a neurons from the HD cortex is reduced in the sFANSseq data. CAP100 scores were calculated as described in Warner et al. (Warner et al., 2022) to represent the cumulative effect of the inherited mutant CAG tract length over the lifetime of the individual.

### sFANS enriched snRNAseq confirms selective vulnerability of *HTR2C* expressing Layer 5a neurons

The finding that a single deep layer pyramidal cell type is missing in the sFANSseq data from HD cerebral cortex, at least early in HD progression, is interesting given the cortical thinning that has been observed in neuroanatomical studies (Reiner and Deng, 2018; Vonsattel et al., 2011; Vonsattel et al., 1985). To explore further the fate of neuronal populations in the HD cortex, we employed snRNAseq (Macosko et al., 2015). Given the relatively low abundance of the Layer 5a *HTR2C* nuclei recovered from the HD donor samples, and to allow us to focus on specific populations in subsequent studies, we used sFANS to subdivide cortical nuclei into four populations prior to snRNAseq: NeuN negative; NeuN positive, SATB2 negative; NeuN positive, SATB2 medium; and NeuN positive, SATB2 high (Figure 3A). To align our sFANSseq data with the snRNAseq data, we used the sFANS data to identify clusters from the snRNAseq. We first generated specificity indices (Dougherty et al., 2010) for all isolated cell-types (Supplementary Figure S3A). Next, we utilized the cell-type specific gene ranking from the specificity index analysis to produce correlation matrices with the snRNAseq clusters (Supplementary Figure S3B) and confirmed assignments based on known marker genes (Supplementary Figure S3C). Cell-types assigned to clusters based on Specificity Index correlations and Uniform Manifold Approximation and Projection (UMAP) were color coded based on these analyses (Figure 3B) identifying the 15 cell classes studied in the sFANS approach in both, HD and control samples. As expected, since specific cell types were targeted in our sFANS analyses, there were additional clusters evident in our snRNAseq data. These additional clusters could be assigned based on known marker gene expression, although three clusters remained unassigned. Lastly, previously published human motor cortex datasets (Bakken et al., 2021) were projected onto our sFANS derived snRNAseq datasets and heatmap correlation matrices were produced to resolve the last remaining cluster identities and to confirm our cluster assignments (Supplementary Figure S3D).

**Figure 3.**
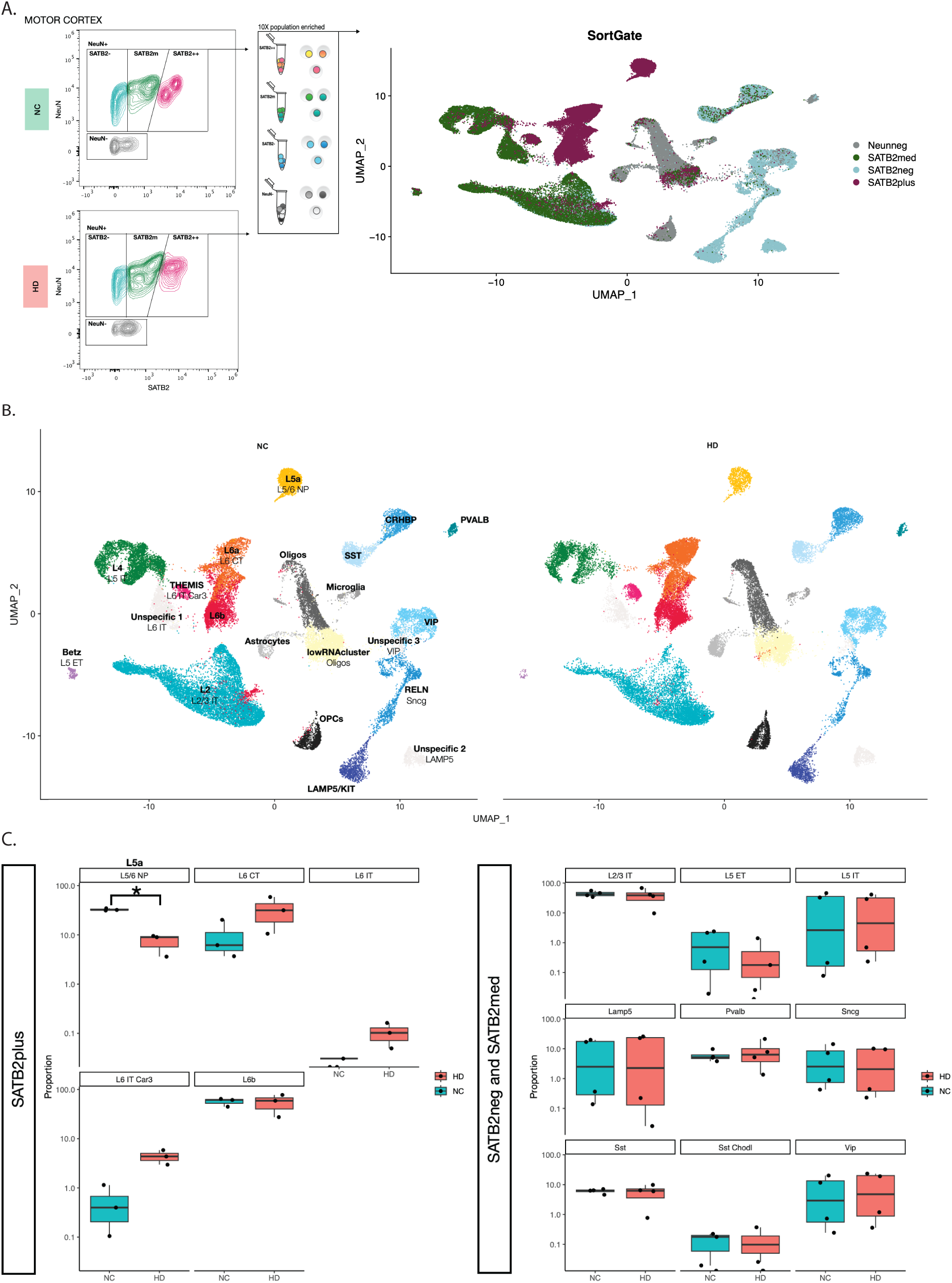
sFANS enriched snRNAseq confirms selective vulnerability of *HTR2C* expressing Layer 5a neurons. (**A**) Schematic representation of sFANS pre-sorting strategies and resulting Uniform Manifold Approximation and Projection (UMAP) plots of the single nuclei RNA sequencing (snRNAseq) data, color coded by sort population (NeuNnegative in gray, NeuNpositive/SATB2negative in light blue, NeuNpositive/SATB2medium in green, and NeuNpositive/SATB2high in dark red). (**B**) Separated UMAPs for NC and HD samples, showing data from full donor datasets for which NeuNnegative, NeuNpositive/SATB2negative, NeuNpositive/SATB2medium, and NeuN positive/SATB2high nuclei could be harvested (number of donors NC=2, HD=2). Cluster assignments in bold and color coding based on sFANS-guided cell-type designations and marker gene expression. Cluster designations in normal font correspond to the correlated clusters in the publicly available human motor cortex datasets (Bakken et al., 2021). (**C**) Quantitation of fraction nuclei in NC vs HD cell-type specific clusters as designated in (Bakken et al., 2021) for NeuNpositive/SATB2negative (number of samples NC=2, HD=2) and NeuNpositive/SATB2medium (number of samples NC=2, HD=2) combined, as well as for NeuN positive/SATB2high (number of samples NC=3, HD=3 with one replicate sample from each group) nuclei (number of donors NC=2, HD=2), reveal statistically significant loss of L5a (L5/6 NP) nuclei from the NeuNpositive/SATB2high populations in HD (adjusted P value < 0.05 in Bonferroni multiple comparison corrected T-test, marked with the asterisk).

To assess HD-dependent differences in abundance of specific cell-types, we analyzed sFANS- enriched snRNAseq data from a total of 18 sorted samples from motor cortex of 2 control and 3 HD donors. These samples were enriched for NeuN negative (n=4, 2 NC, 2 HD), SATB2 negative (n=4, 2 NC, 2 HD), SATB2 medium (n=4, 2 NC, 2 HD), and SATB2 plus (n=6, 3 NC, 3 HD) populations. Comparative analysis of the snRNAseq data from control and HD donors, and quantitation of the number of nuclei recovered for each cluster, revealed several interesting features. First, and most importantly, a clear drop in the number of *HTR2C* expressing Layer 5a nuclei was evident (Figure 3C, marked with asterisk). Second, no loss of nuclei was revealed for any other snRNAseq cluster, including the unassigned clusters not evident in the sFANS data. Third, no new clusters appeared in HD samples in which loss of Layer 5a *HTR2C* expressing nuclei was clearly evident. These data confirm the finding that *HTR2C* expressing Layer 5a neurons are consistently vulnerable in the HD motor cortex, and they demonstrate that this is frank cell loss because we do not recover them in other snRNAseq clusters. These data also suggest that presorting for the nuclei expressing high levels of *SATB2* may be an efficient strategy for focused snRNAseq analyses of these relatively rare deep layer neuronal populations (on average 1.5% out of all NeuN positive nuclei).

It is important to note that the data we have presented in this study precludes assessment of neuronal loss that is variable depending on the clinical presentation or cortical region. For example, Thu et al (Thu et al., 2010) report that loss of calbindin positive interneurons is evident in the motor cortex of HD patients presenting with predominantly motor changes. Loss of these neurons is not observed in the motor cortex of patients presenting with mood and cognitive changes and was not evident when all HD cases were considered together. Although in this study we have analyzed samples from the motor cortex for 13 different HD donors, clinical information regarding their presentation at the time of diagnosis is not available. Furthermore, methodological differences between nuclear counting using sFANS and snRNAseq and antigen staining by conventional immunohistochemistry preclude direct comparisons of these data. For example, calbindin is expressed in at least three populations of human cortical interneurons (<u>Allen Brain</u> <u>Atlas</u>) and SMI32 labels a minority of pyramidal cells in all layers (Del Rio and DeFelipe, 1994; Macdonald and Halliday, 2002). If the decrease (Mehrabi et al., 2016; Nana et al., 2014; Thu et al., 2010) of these antigens in HD motor cortex due to compromised expression due to the disease, or if the fraction of nuclei impacted in any one molecularly defined subclass is small and variable, then a statistically significant difference between control and HD samples as measured by decrease in nuclear number in the motor cortex would not be evident. Similar considerations apply for the more limited data we have collected from other areas of the cerebral cortex. Given the very interesting findings that vulnerable cortical interneuron classes vary based on clinical classification and cortical area, the methodological considerations noted above, and our data demonstrating intermediate levels of CAG expansion in interneurons and upper layer pyramidal cells from HD donors (see below), it will be interesting to extend these studies using a more targeted approach in clinically well characterized cases from early stage HD donors.

### Extensive somatic expansion of the *mHTT* CAG repeat in both vulnerable and resilient cortical pyramidal cell populations

To determine the degree of CAG expansion of *mHTT* in human cortical neurons and its relationship to selective vulnerability, we isolated genomic DNA from each sFANSseq cell population and measured the CAG tract from these populations using Illumina-sequencing across *HTT* exon 1 (Ciosi et al.). Data from a single donor for all analyzed cell types, with the inherited CAG size as defined through mapping of read length-distribution in CAG-sizing data from non-expanding cell types (for example microglia and astrocytes) (red arrow), is shown in Figure 4A. These data reveal very little CAG expansions in the four glial cell types studied (astrocytes, microglia, oligodendrocytes, and Oligodendrocyte Precursor Cells (OPCs)), moderate expansions in each of the four interneuron populations (VIP, RELN, LAMP5, and PVALB), and pronounced expansions in the layer 5a vulnerable neurons, as well as Betz cells, layer 6a and layer 6b pyramidal cell populations that are resilient in HD motor cortex. Representative plots from the motor cortex of four additional donors for the maximally expanding deep layer populations, including Betz cells, confirmed this result (Figure 4B). To extend this analysis to additional cortical regions, CAG expansions were measured in the motor, visual, prefrontal, cingulate and insular cortices of additional HD donor samples. Radar plots of these data from each cell type in each donor display the percentage of nuclei where the *mHTT* CAG tract has expanded beyond the modal inherited CAG length by 3 to 5 repeats (grey), 6 to 20 repeats (black), or by more than 20 repeat units (red). These data also document that somatic expansions occur in nearly all deep layer neurons and that, in approximately half of the deep layer neuron samples analyzed, large expansions were very frequent such that the majority of *mHTT* copies reached CAG tract lengths that carry more than 20 additional CAG units (Figure 4C, see also Supplementary Figure S4 and Table 1 for cell-type specific statistics). These data demonstrate expansions of the *mHTT* CAG repeat in many cell types across the human cerebral cortex, with extensive expansion occurring in both vulnerable and resilient deep layer neurons. It is noteworthy also that the histograms of *mHTT* CAG tract size-distribution of expanded *mHTT* alleles in deep layer neurons appears less uniform in cortical neurons than MSNs (Mätlik et al., 2023). These data demonstrate that somatic CAG expansions are not restricted to vulnerable cell populations in the cerebral cortex. However, since the maximum number of CAG repeats detectable with the MiSeq assay is 113 (Ciosi et al., 2019), further analysis will be required to determine whether the infrequent very long repeats that have been reported in the HD cerebral cortex and striatum (Kennedy et al., 2003) differ between cortical cell types.

**Figure 4.**
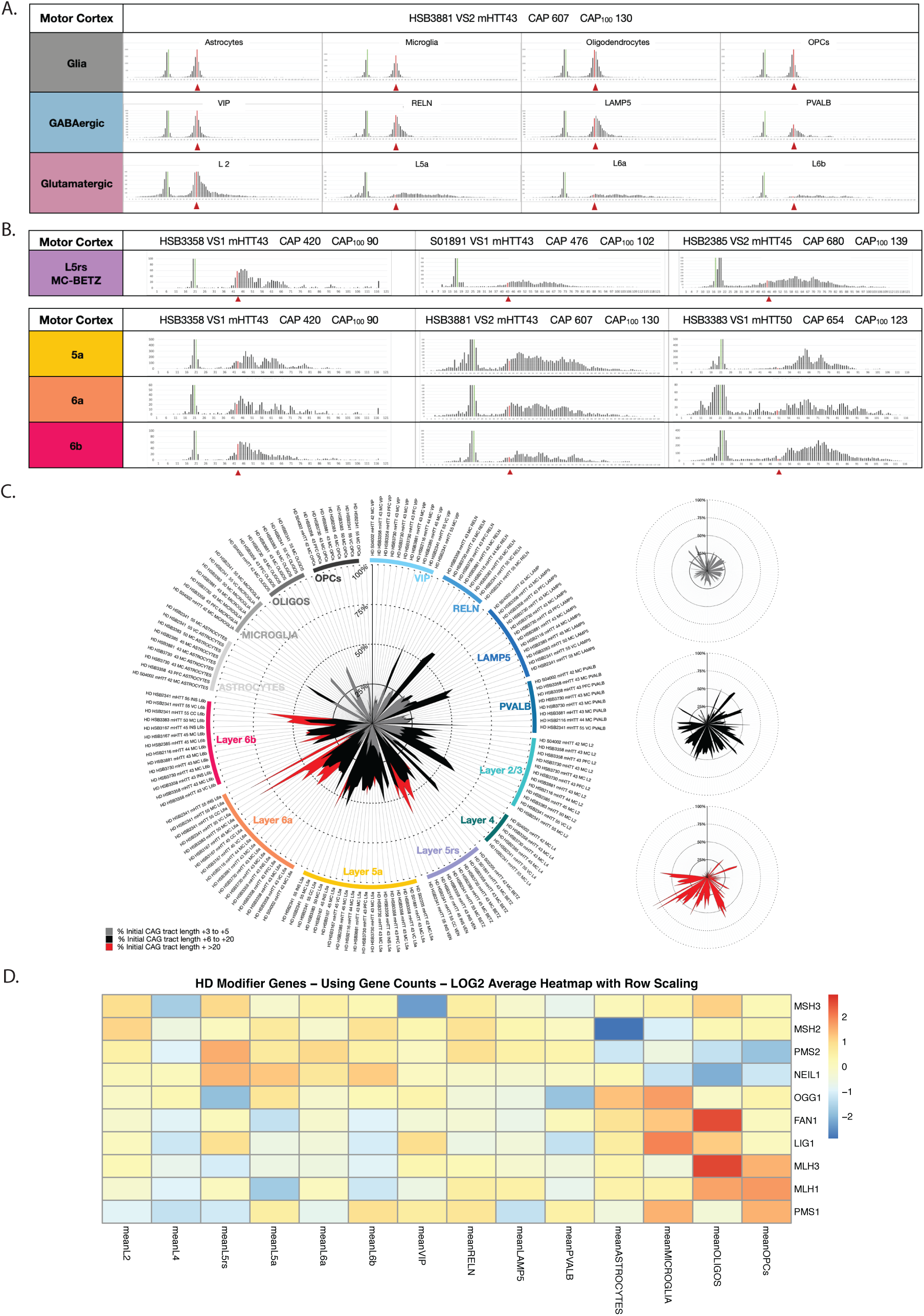
Somatic expansion of the *mHTT* CAG repeat in cortical cell types. (**A**) Representative plots showing CAG length distributions in all cortical cell-types analyzed from one HD motor cortex donor. Red arrow/line indicates the modal allele length defined for each donor through genotyping and assessment of non-expanding glial data from the same donor. (**B**) Representative plots of CAG length distributions in deep layer excitatory neuron populations (L5rs (Betz cells in motor cortex), layer 5a (L5a), layer 6a (L6a), layer 6b (L6b)) from five HD donors. (**C**) Radar plot showing data from all cell-types of eleven HD donors, and all cortical regions analyzed. Data are color coded by length of CAG expansions greater than the modal length in striatal glia, with 3 to 5 repeats in grey, 6 to 20 repeats in black, and over 20 repeats in red. (**D**) Heatmap shows log-2 transformed average DESeq-normalized expression levels of selected HD modifying genes across all cell-types at baseline, utilizing data from six control donors.

**Table 1.**

| **MOTOR**<br>**Cortex** | **CAG expansion beyond inherited mutant**<br>**allele length** |  |
| --- | --- | --- |
|  | **Mean** | **SD** |
| **L2** | 11.2 | 3.0 |
| **L4** | 9.6 | 4.8 |
| **BETZ** | 18.9 | 4.1 |
| **L5a** | 16.4 | 3.3 |
| **L6a** | 21.6 | 4.0 |
| **L6b** | 17.3 | 5.3 |
| **VIP** | 5.0 | 2.9 |
| **RELN** | 8.4 | 5.9 |
| **LAMP** | 9.0 | 5.6 |
| **PVALB** | 6.8 | 1.8 |
| **ASTROCYTES** | 3.7 | 1.9 |
| **MICROGLIA** | 2.6 | 0.8 |
| **OLIGOS** | 3.5 | 0.7 |
| **OPCs** | 2.8 | 1.0 |

### The vulnerable layer 5a *HTR2C* expressing cells in primates are corticostriatal projection neurons

Excitatory neurons in the cerebral cortex are primarily pyramidal cells (Cajal, 1999) that send projections to a variety of brain structures and targets (Cajal, 1999; Elam et al., 2021; Usrey and Sherman, 2019). Layer 5 neurons project to subcortical targets, and detailed studies in the mouse cortex have demonstrated that they have distinct physiological properties (Groh et al., 2010; Hattox and Nelson, 2007). Layer 6 neurons generally either project subcortically to target nuclei in the thalamus or have axons that ascend within the cortex (Baker et al., 2018). Since a variety of human imaging studies of HD have documented alterations in the connectivity of multiple areas of the cortex with the striatum, we investigated next the projection targets of the *HTR2C* expressing layer 5a neurons in the primate cortex. To do so, we took advantage of the very detailed mapping of striatal projecting neurons in the adult rhesus macaque previously reported by Weiss et al (Weiss et al., 2020), employed a retrograde AAV2 tracer virus expressing eGFP (AAV2.retro) to transduce neurons in the caudate nucleus and putamen. Immunohistochemical detection of the eGFP gene delivered by the viral vector revealed many cortical and subcortical neurons that project to the striatum in the non-human primate (NHP) brain. The highest densities of retrogradely transported virus were detected in anterior regions of the cerebral cortex, including the prefrontal and motor areas (Weiss et al., 2020). To map *HTR2C* expressing cells in the AAV2.retro labeled macaque brains, sections from these animals were examined and sub-dissected to use for analysis of the motor and cingulate cortex. Immunofluorescence data collected from these sections revealed robust staining of eGFP in many layer 5 pyramidal neurons (Figure 5A). To simultaneously analyze expression of both eGFP and *HTR2C*, we used RNA scope *in situ* hybridization (ISH) (Wang et al., 2012). Dense signal for eGFP (green) in layer 5 neurons using this assay was evident even in low power images (Figure 5B), as well as less robust signal from the *HTR2C* probe (red). At higher magnification, specific eGFP expression from the AAV2.retro virus as well as endogenous macaque *HTR2C* expression could be evaluated as the increased grain counts within and surrounding DAPI positive nuclei relative to background for each probe (Figure 5C). Sections stained with both RNAscope probes and DAPI were scanned at 20X magnification, tiled, and analyzed using HALO (Erben and Buonanno, 2019). To determine whether corticostriatal cells expressing eGFP from AAV2.retro also express *HTR2C*, parameters were set to reflect three classes of colocalization signal: cells strongly positive for eGFP, cells strongly positive for both eGFP and *HTR2C*, and cells strongly positive for eGFP and weakly positive for *HTR2C* (Figure 5B). Representative data for single sections from cingulate and motor cortex reveal that 41% (motor cortex) and 32% (cingulate cortex) of the eGFP expressing neurons are positive for high levels of *HTR2C*, with an additional 67% (motor cortex) and 68% (cingulate cortex) weakly positive in cingulate and motor cortex respectively (Figure 5D). These data demonstrate that macaque corticostriatal projection neurons express *HTR2C*, the marker gene specific for the vulnerable neuron type in HD.

**Figure 5.**
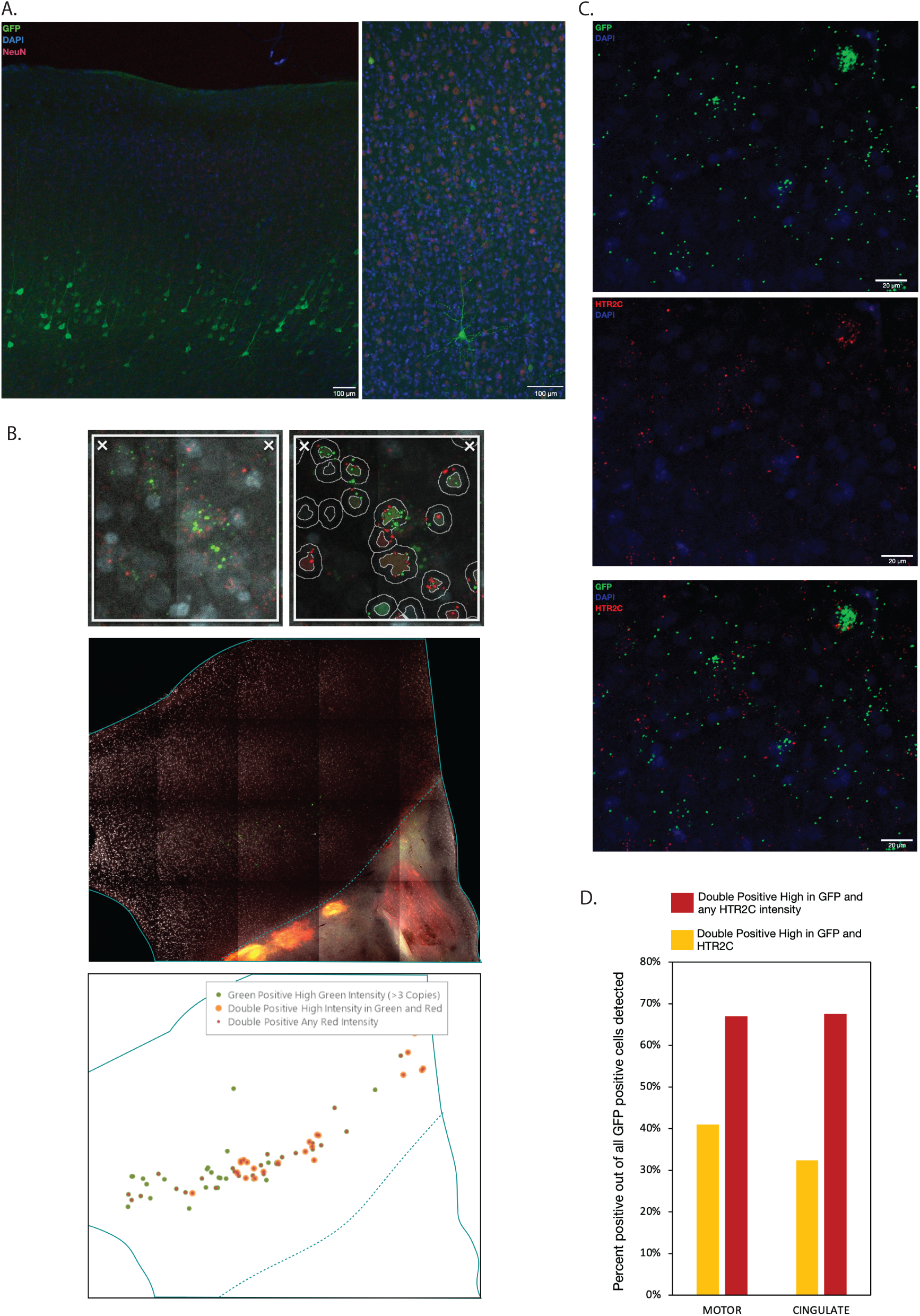
Vulnerable layer 5a *HTR2C* expressing cells in primates are corticostriatal projection neurons. (**A**) Immunohistochemistry (IHC) experiment results from macaque cortex confirm eGFP positive staining of layer 5 pyramidal neurons. (**B**) Representative in situ hybridization (ISH) data analyses of NHP cingulate cortex, using HALO software for distribution analyses. Delineation of DAPI positive nuclei and eGFP as well as *HTR2C* RNA signal puncta quantitation, using HALO analyses tool is shown in higher magnification in top panels. Analyses results, revealing eGFP and *HTR2C* positive signal in deep layer populations at low magnification (middle and bottom panels). (**C**) High magnification images from the cingulate cortex show positive signal for eGFP (green) and *HTR2C* (red). (**D**) HALO quantification results from single sections from cingulate and motor cortex reveal 41% (motor cortex) and 32% (cingulate cortex) of double positive for eGFP and high *HTR2C* neurons and 67% (motor cortex) and 68% (cingulate cortex) neurons being positive for eGFP and weakly positive for *HTR2C*.

It is noteworthy that the data we have collected in this study are consistent with both previous ISH studies of human *HTR2C* expression (Supplementary Figure S5B, brain-map.org/ish/HTR2C) and regional variation in retrogradely labeled cells in the macaque cerebral cortex (Weiss et al., 2020). Thus, the decline in the number of retrogradely labeled neurons noted as one moves from anterior to posterior regions in the macaque cortex is also reflected in the human *HTR2C* ISH data from the frontal, temporal and visual cortex in the Allen Brain Atlas (brain-map.org), and in the differences in the recovery of *HTR2C* expressing layer 5 neurons in the sFANS studies of samples from the prefrontal, motor and visual cortex (Supplementary Figure S5C). For example, although the precise location of the samples used for the sFANS from these areas is difficult to assess, very small numbers of *HTR2C* expressing nuclei were recovered from samples identified as primary visual cortex. Furthermore, in the L5a nuclei recovered from the visual cortex, less somatic expansion is apparent (Supplementary Figure S4).

### Transcriptional responses in the HD cerebral cortex

The fact that the somatic expansion occurs to a similar degree in all deep layer projection neuron classes we have analyzed yet, while it is the Layer 5a corticostriatal neurons that are selectively lost in HD is of high interest. To gain insight into the impact of *mHTT* with long CAG tracts on cellular functions in each of these cell types, we compared their transcriptional responses to *mHTT*. Cell-type specific differential gene expression (DESeq2) analyses revealed over 1,000 differentially expressed genes in layer 5a, layer 6a, and layer 6b neurons from HD versus control donor samples. Analyses of layer 5rs neurons revealed less than 300 differentially expressed genes (Figure 6A). MA plots show log2FC and basemean of differentially expressed genes between HD and control donor samples in black, with significantly upregulated genes in red, and significantly downregulated genes in blue for layer 5a, layer 6a, and layer 6b (Figure 6B). Principal component analyses (PCA) of the three deep layer neuron populations, including layer 5a, layer 6a, and layer 6b neurons, revealed separation of samples based on sex (PC1) and disease status (PC2) (Supplementary Figure S6A). Samples from layer 5rs neurons however, showed no clear separation based on disease status, with the exception of a weak trend for separation in PC3 of Betz cells (Supplementary Figure S6C). Analyses of layer 5rs samples confirmed no clear pseudotime trajectory or trend according to disease status (Supplementary Figure S6C). Pseudotime analyses, using our sFANSeq data from the deep layer cell-types further confirmed a clear progression trajectory of molecular alterations with increasing CAP scores (Figure 6D) for layer 5a, layer 6a, and layer 6b neurons.

**Figure 6.**
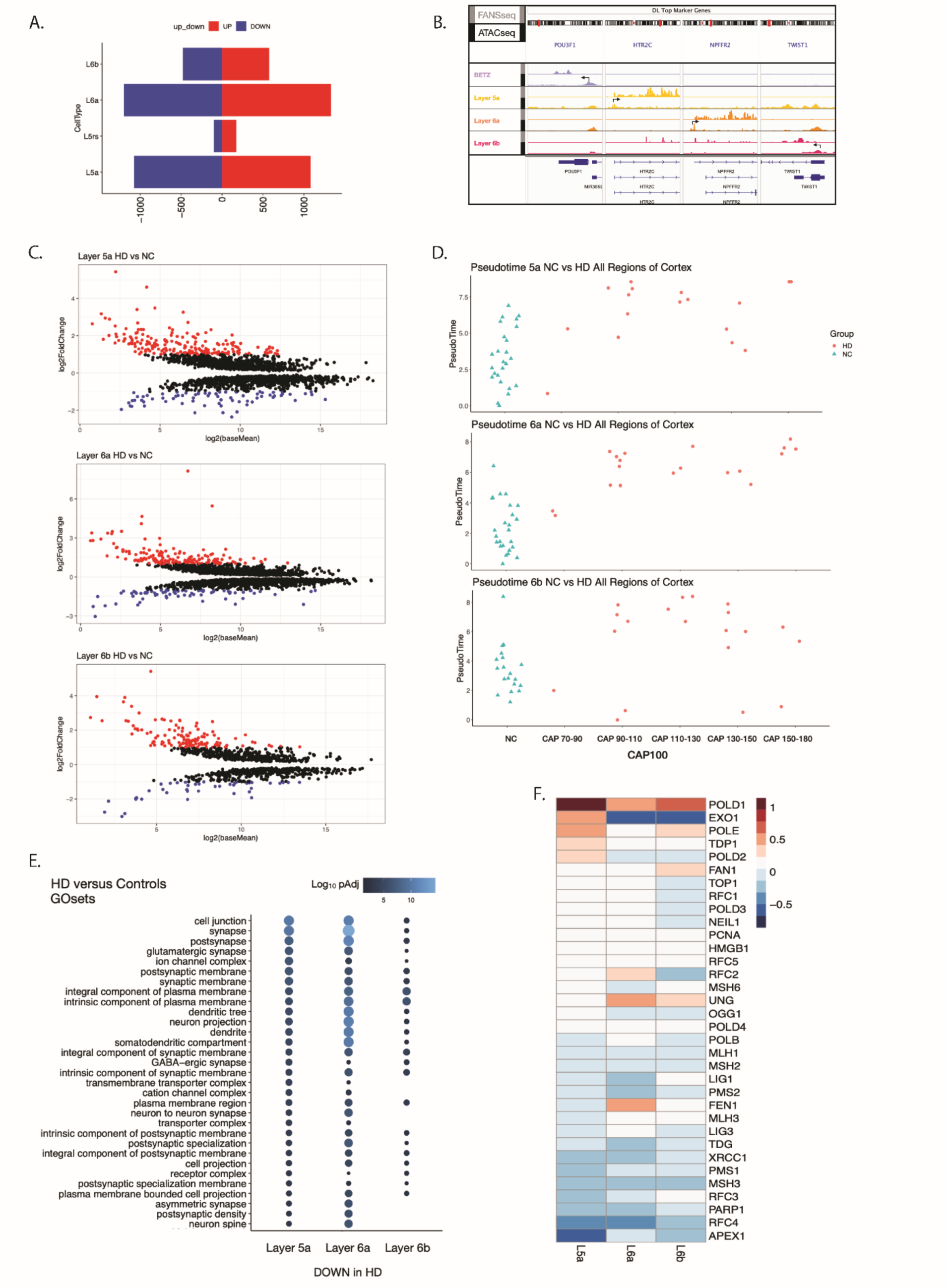
Transcriptional responses in the HD cerebral cortex. (**A**) Number of differentially expressed genes (adjusted P value < 0.05 by DESeq2) in the comparison of datasets from HD donors versus controls and after gene list filtering, based on the detection of accessible promoters using ATACseq. (**B**) Representative distribution of sFANSeq and ATACseq reads from motor cortex mapped to deep layer cell-type specific marker genes. (**C**) MA plots illustrating log2FC and log2baseMean of differentially expressed genes in layer 5a, layer 6a, and layer 6b samples from HD versus NC. Significantly up regulated genes are plotted in red, significantly down regulated genes are plotted in blue. Non-significant but differentially expressed genes are plotted in black. (**D**) Bulk pseudotime analyses results for layer 5a, layer 6a, and layer 6b, depicting molecular responses along the pseudotime axis in control and in HD samples across CAP scores. (**E**) Results from Gene Ontology Cellular Component pathway enrichment analyses of genes downregulated in HD versus controls (p adj < 0.05 by DESeq2 and filtered for genes with accessible promotors detected by ATACseq) in deep layer excitatory neuron populations, including layer 5a, layer 6a, and layer 6b nuclei. (**F**) Differential gene expression between HD and control samples from layer 5a, layer 6a, and layer 6b are shown for a set of genes of interest.

To reveal gene expression programs altered in the deep layer neuron cell classes that showed molecular alterations based on disease status, we performed gene ontology (GO) analyses. To this end, we first filtered the cell-type specific differential gene expression lists from our DESeq2 analyses based on the cell-type specific accessible promotor peaks revealed through ATACseq (Figure 6B). This step allowed us to focus our gene lists for the GO analyses to those genes that were truly accessible in the specific cell-types. As shown in Figure 6E, expression of genes involved in dendritic and synaptic functions are decreased in layer 5a, layer 6a, and layer 6b neurons that undergo extensive somatic expansion of the *mHTT* CAG tract in HD. These data are generally consistent with studies in mouse models of HD (Plotkin and Surmeir, 2015; Veldman and Yang 2018), and with human imaging studies indicating disconnection between the cortex and striatum (Espinoza et al., 2019; Unschuld et al., 2012), and they support the hypothesis that *mHTT* carrying long polyglutamine tracts can disrupt many aspects of dendritic and synaptic function. Analyses of differential expression of genes of interest between HD and controls revealed multiple DNA repair genes as differentially expressed between HD and controls (Figure 6F). Analyses of POLD1 expression in layer 5a, layer 6a, and layer 6b control donor samples revealed no gradient across the six cortical regions (Supplementary Figure S6B).

To provide further insight into the increased vulnerability of corticostriatal pyramidal cells in HD, we compared the transcriptional responses of layer 5a *HTR2C* neurons to layer 6a and layer 6b pyramidal cells that have equivalent levels of CAG expansions in HD but remain relatively resilient against cell death (Supplementary Figure S6D). In general, these data revealed similar pathways as those discussed above: depressed expression of pathways involved in neuronal membrane functions required for support of dendritic and synaptic functions. Pathways that show enrichment of upregulated gene sets in layer 5a neurons include those involved in membrane flux and delivery of cell surface proteins to their sites in the membrane. These data indicate that altered synaptic function of L5a and, perhaps, L6 deep layer neurons may result in their dysfunction, and they suggest that altered corticostriatal connectivity may play an important role in the selective loss of L5a corticostriatal pyramidal cells in HD.

## DISCUSSION

Although degeneration of the striatum and the associated disturbances of motor control are the most visible features of Huntington’s disease, cognitive decline and psychiatric symptoms occur early in HD and they are accompanied by functional and anatomical alterations in the cerebral cortex (Hickman et al., 2022; Mehrabi et al., 2016; Nana et al., 2014; Nopoulus, 2010). Cortical dysfunction in HD is thought to be the result of variations in the vulnerability of specific cell-types that depends both on the clinical presentation of the case studied and the cortical region analyzed (Kim et al., 2014; Mehrabi et al., 2016; Thu et al., 2010). Here we have used serial fluorescence activated nuclear sorting (sFANS) for comprehensive molecular profiling to explore pathogenesis in six human cortical regions, with a focus on the motor cortex, in Huntington’s disease. Our data identify Layer 5a *HTR2C* expressing corticostriatal pyramidal cells as consistently vulnerable in the HD cerebral cortex, they establish that extensive somatic expansion of the *mHTT*-CAG tract occurs in both vulnerable L5a pyramidal cells and in their more resilient L6a, L6b, and L5rs projection neuron neighbors, and they demonstrate that transcriptional programs disturbed in the HD cerebral cortex are enriched in genes encoding synaptic proteins. Taken together, these findings suggest that both somatic expansion of the *mHTT* CAG tract and disruption of corticostriatal connectivity may contribute to selective vulnerability in HD. It is notable also that the transcriptional disturbances evident in deep layer neurons in the cerebral cortex are quite different than those we have observed in the human striatum (Mätlik et al., 2023), suggesting that the pathways influencing selective vulnerability in the cerebral cortex and striatum in Huntington’s disease are distinct.

### Serial fluorescence activated nuclear sorting (sFANS)

Studies of human tissue are challenged by the paucity of appropriate samples for analysis of early phases of disease. Given the many cell types present in the human cerebral cortex (Cajal, 1999), the advantages of population based molecular studies evident in our studies of the human cerebellum (Xu et al., 2018), the desire to use a tissue sparing strategy for studies of HD donor samples, we developed sFANS for deep molecular analysis of all major cortical cell classes from a single tissue sample. The approach first involved sorting of nuclei into four pools based on the expression of *NEUN* and *SATB2*, followed by a second sort for each of these pools into subpopulations of nuclei based on the presence of *BCL11b*, *TLE4*, *EAAT1*, *IRF8* or *OLIG2*. Although most of these markers do not identify a single cell type, the multiplexed nature of the sort and quantitative differences in expression resulted in purification of nuclei for each of the major excitatory, inhibitory, and glial cell classes in the human cortex, as well as region specific cell types such as Betz and von Economo neurons. Importantly, the yield from the sFANS strategy was sufficient to collect thousands of nuclei from each cell type for analysis of *mHTT* CAG DNA expansion, epigenetic, and transcriptional data for each class of nuclei. This enables accurate identification of active gene expression through cross-correlation of the RNAseq data with promoter accessibility as revealed by ATACseq (Mätlik et al., 2023), and removal of ambient RNA reads that have been recognized as an important confounding factor in snRNAseq (Caglayan E., 2022). Furthermore, although not used for these studies, probes against individual cell types can allow targeting of a single, previously identified cell type for analysis if required (Supplementary Figure S2C). Finally, as demonstrated here, distortions in the staining patterns observed on sFANS plots from HD donors can provide a preliminary readout of the relative abundance of each population in comparisons of cortical regions of donor samples. It is instructive, for example, that the differences visible on sFANS plots for deep layer neuron nuclei were the first indications of cell type specific HD pathology to emerge in this study.

### Loss of layer 5a *HTR2C* expressing corticostriatal pyramidal cells in Huntington’s disease

The data we have presented here strongly support the conclusion that the layer 5a *HTR2C* cells that are lost in the human cerebral cortex early in HD are pyramidal cells that project to the striatum. Although we have analyzed a relatively small number of HD donor cases (N=13), loss of *HTR2C* expressing cells is documented by both sFANSseq and snRNAseq across multiple brain regions and in many samples. However, due to the lack of clinical information available for the donors we have studied and methodological considerations, our data cannot be used to address previous findings demonstrating variable loss of interneurons and pyramidal cell subclasses that depends on both clinical presentation and cortical area (Hedreen et al., 1991; Hickman et al., 2022; Kim et al., 2017; Mehrabi et al., 2016; Nana et al., 2014; Nopoulus, 2010; Thu et al., 2010). Although extension of these studies to additional, well defined HD cases and many cortical regions will be required to understand the complexity of selective neuronal loss during disease progression, our data establish that L5a pyramidal cells are consistently vulnerable in the cerebral cortex in HD. While it remains possible that the *HTR2C* neurons in the human cortex are not homologous to the corticostriatal projection neurons that were labeled by *HTR2C* and AAV2.retro in the macaque cortex, we view this as unlikely because of the abundance gradient of these cells along the antero-posterior axis that is common to the two primate species (Bernard et al., 2012; Weiss et al., 2020).

The selective vulnerability of corticostriatal neurons in HD that we have documented here relative to the more resilient layer 6 neurons and Betz cells is interesting given the hypothesis that these cells would be preferentially impacted in HD patients (McColgan et al, 2020), and with respect to the functional imaging studies of *mHTT* carriers (Espinoza et al., 2019; Unschuld et al., 2012). Corticostriatal pyramidal cells are intratelencephalic (IT) neurons that can project bilaterally through the corpus collosum to the contralateral cortex and striatum (Shepherd, 2013, Baker et al, 2018, Hooks et al, 2018). The loss of L5a pyramidal neurons documented here raises the interesting possibility that their early loss could contribute to the callosal atrophy observed in premanifest and early HD (Tabrizi et al, 2011; Crawford et al, 2013). Furthermore, since these cells are direct presynaptic partners of striatal MSNs, an understanding of the relative timing of their dysfunction and loss of these two cell-types in HD is essential. Longitudinal studies of the functional and connectivity changes in premanifest and symptomatic HD subjects imaging studies (Georgiou-Karistianis et al., 2013; Wolf et al., 2011), as well as PET (Kuwert et al., 1990), have suggested also that the cortical activation in pre-manifest HD may reflect a compensatory response to decreased functional connectivity between the cerebral cortex and striatum as an early sign of progression, and that the diminished activation seen in symptomatic stages may reflect the inability of the brain to continue to compensate as neuronal damage accumulates. This places an important emphasis on delineating changes in the properties of these cells in HD (see below), and in the timing of their dysfunction and loss across cortical regions. Furthermore, the robust expression of *HTR2C* in both corticostriatal pyramidal cells and striatal projection neurons suggests that further development of the *HTR2C* imaging probes (Kim et al., 2017) for use in longitudinal studies could be particularly important for understanding the relationships between corticostriatal disconnection and neurodegeneration in HD.

### Widespread somatic expansion of *mHTT* CAG repeats in the HD cerebral cortex

In the striatum, somatic expansion of the *mHTT* CAG tract is evident exclusively in striatal projection neurons and cholinergic interneurons expressing choline acetyltransferase (CHAT+ INs) (Mätlik et al., 2023). Significant CAG expansions are rare in other striatal interneuron populations and glia. While the very limited expansion in glia observed in striatum is also evident in cerebral cortex, all classes of cortical neurons we have analyzed display somatic instability of the *mHTT* CAG tract. A significant fraction of cortical interneurons, for example, have tracts expanded by more than 5 additional repeat units. This is in contrast to the lack of somatic CAG expansions in many striatal interneurons, including those expressing *SST*, *PVALB*, and *TAC3* (Mätlik et al., 2023). Interestingly, striatal and cortical interneurons have a common developmental origin within the medial ganglionic eminence as progenitors of both (Knowles et al., 2021) span the cerebral cortex and striatum (Reid and Walsh, 2002). One might argue, for example, that this provides evidence that the differential CAG expansion levels in related interneuron cell types, as evident in our data, may arise late in development as cortical and striatal interneurons diverge as a consequence of local cell-cell interactions in the cortex.

The most extensive CAG expansions in the HD cortex are evident in the four (L5rs, L5a, L6a, and L6b) deep layer pyramidal cell populations we have analyzed (Figure 4B). These data demonstrate expansion in the majority of neurons in these cell types to as many as 100 *mHTT* CAG repeats does not differ between the cortical cell types that are vulnerable (layer 5a corticostriatal neurons) and resilient to degeneration (Betz neurons and layer 6 populations). However, given the upper limit of the MiSeq assay to detection of 113 repeat units, of our data do not address possible differences in the fraction of nuclei carrying very long repeats (Kennedy et al., 2003) between vulnerable and resilient cell types that have been proposed as particularly important in models of HD cell loss (Kaplan et al., 2007).

### Cell-type specific patterns of somatic CAG instability

Somatic CAG instability is influenced by specific features of the locus that is expanding, its transcriptional and epigenetic state, and trans-acting factors that directly act at the repeat locus to enhance or suppress somatic instability (Nakamori et al., 2011) (Barbe and Finkbeiner, 2022). For example, in striatal medium spiny neurons (MSNs), which show the greatest CAG expansions in that structure, the expression of *MSH2* and *MSH3* is higher both at the nuclear transcript and protein level relative to other striatal cell types (Mätlik et al., 2023). One explanation of the expansions in MSNs and stability in striatal interneurons is that MutSβ (MSH2-MSH3), a driver of expansions, can inhibit excision of excess repeats by FAN1, a suppressor of expansions, while at lower MutSβ levels FAN1 can effectively excise the excess repeats (Mätlik et al., 2023). In the deep layer cortical neurons that display the most dramatic expansion (L5a, L6a, L6b, and Betz cells) elevated expression of the genes encoding *MSH2* and *MSH3* is not evident relative to neurons and glia that are relatively stable (Figure 4D). This result may indicate that the rate limiting factors driving somatic expansion may differ between cell types, and we note that the expression of *FAN1* in these neurons is low (Figure 4D). While this supports the general idea that the stoichiometric levels of DNA repair proteins can in part explain brain region- and cell type-specific levels of somatic expansions (Goula et al., 2012; Lopez Castel et al., 2009; Mason et al., 2014; Mätlik et al., 2023; Slean et al., 2016; Tome et al., 2013; Tome et al., 2011), the interactions of repair proteins and the variables affecting instability are complex. For example, it is unclear how the increased *POLD1*, a candidate modifier of HD (Lee et al., 2022), in L5 neurons and the reduced *EXO1* in L6a and L6b neurons, two of the most differentially expressed between HD and controls (Figure 6F), can affect CAG expansions.

It is noteworthy also that the histograms of *mHTT* CAG tract size-distribution of expanded *mHTT* alleles in deep layer neurons appears less uniform in cortical neurons than MSNs (Mätlik et al., 2023). Thus, in all samples measured to date, the profile of CAG tract lengths expansion is more distributed and spread over a wider range of CAG repeat lengths in the cortex than in striatal MSNs. Although systematic analysis of additional donors will be required to validate this finding, it seems likely that this observation reflects differences in the stoichiometry or activity of DNA repair complexes operating at the *mHTT* locus in cortical and striatal neurons. While the data do not lead us to conclude that the rate limiting steps for somatic expansion of the *mHTT* CAG tract differ between cell types, they are consistent with the recent report that variants in known HD modifying candidate genes act differentially on motor and cognitive domains of HD (Estrada-Sanchez and Rebec, 2013; Layburn et al., 2022; Lee et al., 2022; Waldvogel et al., 2015). Further in depth studies of the *mHTT* locus in deep layer cortical neurons, transcription of both *mHTT* and its antisense transcripts, and studies protein complexes present at the locus in different human neuronal cell types will be required to understand cell type specific expansion. We believe that the ability to isolate and characterize large numbers of nuclei from HD donor samples using the sFANSseq approach is important for advancing our knowledge of this process in the human brain.

### *mHTT* toxicity

Given the finding that extensive somatic *mHTT* CAG expansions are evident in both vulnerable and resilient deep layer pyramidal cells in the HD cerebral cortex, we believe that comparative studies of the transcriptional events occurring in these cell types may be particularly informative. The high probability that significant levels of mutant huntingtin accumulates in each of these cell types, which is supported by recent findings of regionally variable but pronounced aggregation in cortical layers 5 and 6 (Hickman et al., 2022), has important functional implications. It seems likely, for example, that these alterations cause significant dysfunction in all the expanding deep layer populations, disrupting any of the cells to which they connect. For example, many of the functional GO categories that are enriched for genes with disease-associated downregulation are involved in HD and reflect dendritic and synaptic functions (Figure 6E). Furthermore, enhanced expression of genes involved in plasma membrane function and maintenance, and assembly of important membrane complexes, suggest homeostatic responses may be engaged in the affected neurons in response to the deficits in mechanisms of synaptic support. These human data support previous studies of synaptic and connectivity defects in HD mouse models that have revealed widespread defects in cortico-striatal communication (MacDonald et al., 2003; Veldman and Yang, 2018), and they are in line with the prediction that white matter changes are associated with altered transcription of synaptic genes in HD (McColgan et al., 2018) and with the suggestion that more extensive deficits in the corticostriatal thalamocortical (CSTC) circuit are likely to occur in HD (Barry et al., 2022). Although our data from the human cerebral cortex and those previously reported in mouse models (MacDonald et al., 2003; Veldman and Yang, 2018) indicate that similar processes are impacted, the gene by gene details of neuronal responses to mutant huntingtin in human and mouse deep layer neurons differ. This is not surprising given the well documented species-specific differences that occur in many neuronal cell types (Hodge et al., 2019; Xu et al., 2018) and the ubiquitous expression of *mHTT* with long polyglutamine tract in HD mouse models, in contrast to our findings of cell-type specific somatic CAG tract expansion in human samples.

### Concluding remarks

The use of sFANS to characterize cell type-specific gene expression profiles and HD-associated expression changes in the cerebral cortex illustrates the applicability of this method to the studies of this brain region in the context of other neurological diseases, including ones where tissue availability can be a limiting factor. The analyses of the molecular events that we have presented here are the beginning of an effort to understand in detail HD pathophysiology at the molecular level in the human cerebral cortex. Additional studies will be required to assess variable loss of specific cell subpopulations in HD (Kim et al., 2014; Mehrabi et al., 2016; Nana et al., 2014; Thu et al., 2010) and to determine whether differences in the frequency of very long CAG repeats in mHTT correlates with the selective vulnerability of L5a neurons (Kaplan et al., 2007; Kennedy et al., 2003). Nevertheless, our data provide evidence that expansion of the *mHTT* CAG tract in the majority of neurons in a given projection neuron subtype is not likely not to be the only mechanism that determines the rate of cell loss in HD cerebral cortex, and they suggest that altered circuit function may contribute to cell loss early in the disease. We believe the depth of the data we have been able to collect by sFANS, coupled with the ability to filter out ambient RNA contamination using promoter accessibility data, will enable many additional comparative studies to gain insight into specific aspects of HD progression and into general questions surrounding selective cellular vulnerability in human disease.

## Acknowledgements

This study was supported by funding from the CHDI Foundation, and we are thankful to Dr. Thomas F. Vogt and Dr. Jian Chen for helpful discussions throughout the project. We thank Nicholas Didkovsky, Cuidong Wang and Dr. Kärt Mätlik for advice on coding and data visualization. We thank Dr. Christopher E. Pearson for input and constructive criticism of the manuscript. We are also grateful to the Rockefeller University Genomics Resource Center for advice and support. C.P. Christina Pressl, MD MS is in part supported by the Stavros Niarchos Foundation as the Stavros Niarchos Foundation Scholar. The research was in part supported by grant #UL1TR001866 from the National Center for Advancing Translational Sciences (NCATS), National Institutes of Health (NIH) Clinical and Translational Science Award (CTSA) program. We thank the NIH Neuro Bio Banks, in especially the Miami Brain Endowment Bank as well as Sabina Berretta, director of the Harvard Brain Tissue Resource Center for providing biological samples. We thank all donors and families without whom this work would not be possible.

## Author Contributions

C.P and N.H. conceptualized and designed the study. C.P. and P.D. performed the experiments.

K.M. set up the CAG sizing assay and analysis strategy, optimized TrueBlack quenching, led the efforts on ATACseq-based filtering, and gave crucial guidance and support on a set of visualizations shown in this manuscript. D.D., provided human brain samples. J.MB., W.L., and A.W. provided NHP samples. C.P. analyzed the data in collaboration with J.D.L, M.R.P. and T.C..C.P. and N.H. wrote the manuscript. All authors discussed the results and commented on the manuscript. N.H. acquired research funding.

## SUPPLEMENTAL MATERIALS

List of supplemental files:

Table S1 — Metadata for Tissue Donors.xlsx

Table S2 — Metadata for sFANSseq samples used for analyses.xlsx

Table S3 — ATACfilteredSigDESeq2Genes_HDvsNC_L5aL6aL6b.xlsx

Table S4 — ATACfilteredGOresultsDL_HDvsNC_L5aL6aL6b.xlsx

Table S5 — ATACfilteredGOresults_HDL5avsHDL6.xlsx

## SUPPLEMENTARY FIGURE TITLES AND LEGENDS

**Supplementary Figure 1.**
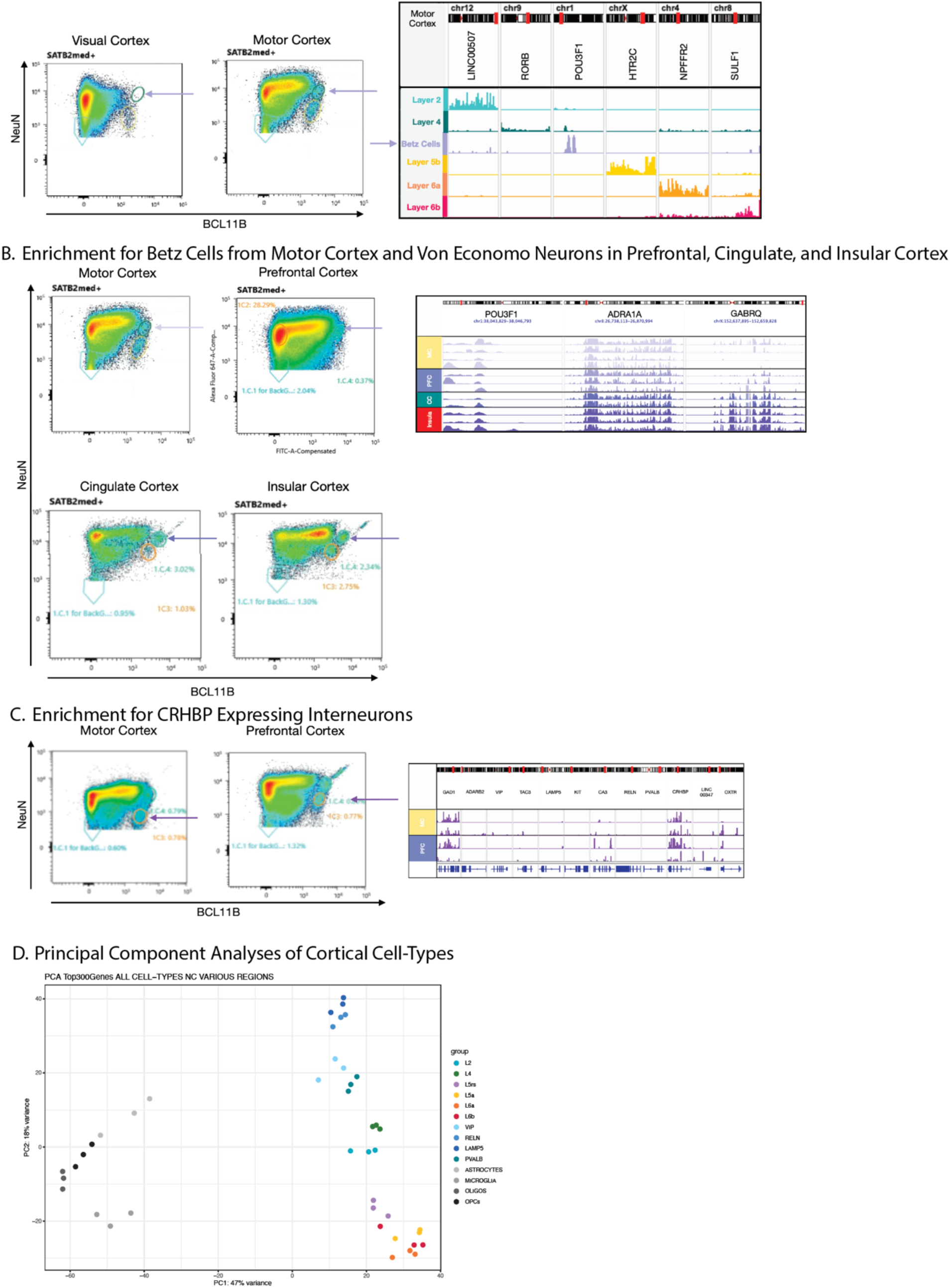
(**A**) sFANS plots show the position of a motor cortex specific population within the SATB2 medium plus range. Resulting marker gene expression profiles from this isolated population (labeled Betz Cells, purple tract) alongside example IGV tracts from layer2, layer 4, layer 5, layer 6a, and layer 6b excitatory cortical neurons, reveal the expression of *POU3F1*. (**B**) sFANS plots from motor, prefrontal, cingulate, and insular cortex show position of isolated populations from the SATB2 medium plus range. Resulting marker gene expression profiles show enrichment for *POU3F1*, *ADRA1A*, and *GABRQ* expressing neurons. (**C**) sFANS plots from motor and prefrontal cortex show the position of an isolated population enriched for the marker gene CRHBP as shown through visualization of resulting RNAseq profiles in IGV. (**D**) Principal Component Analyses of control data examples for layer 2, layer 4, layer 5rs, layer 5a, layer 6a, layer 6b, VIP, RELN, LAMP5, PVALB, Astrocytes, Microglia, Oligodendrocytes, and Oligodendrocyte Precursor Cells (OPCs).

**Supplementary Figure 2.**
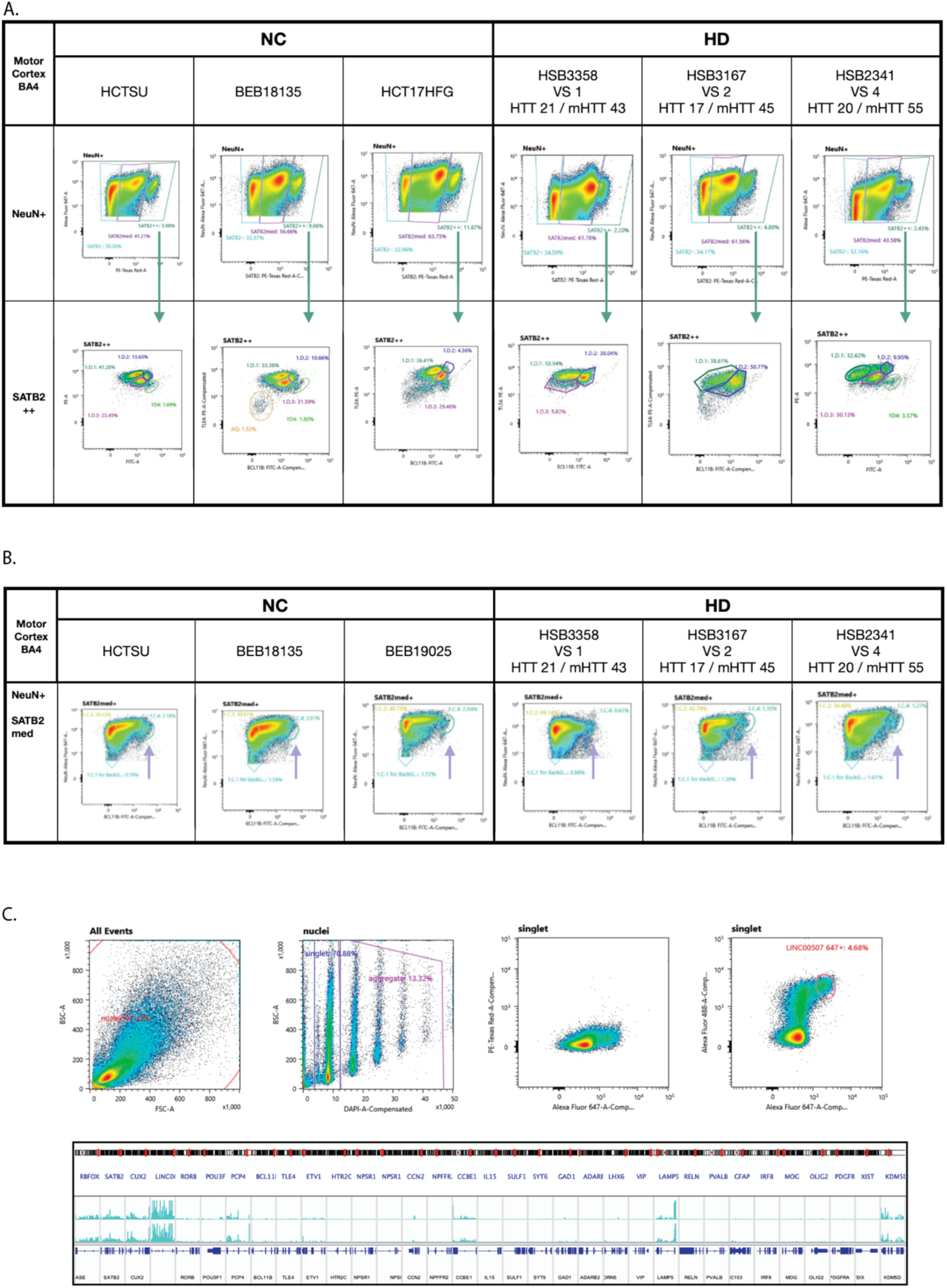
(**A**) sFANS plots showing examples of the distribution of NeuN positive nuclei along the SATB2 negative, SATB2 medium, and SATB2 high range (top row) for three controls and three HD donors. Bottom row shows the distribution of NeuN positive SATB2 high nuclei along the BCL11B and TLE4 ranges, revealing a distortion of the fluorophore signal of these populations in HD. **B**) sFANS plot examples show the position of the Betz cell enriched marker gene population for three controls and three HD donors. (**C**) Example sFANS plot and resulting IGV tracts confirm the isolation of cortical layer 2 neurons using RNA probes against the LINC00507 marker gene.

**Supplementary Figure 3.**
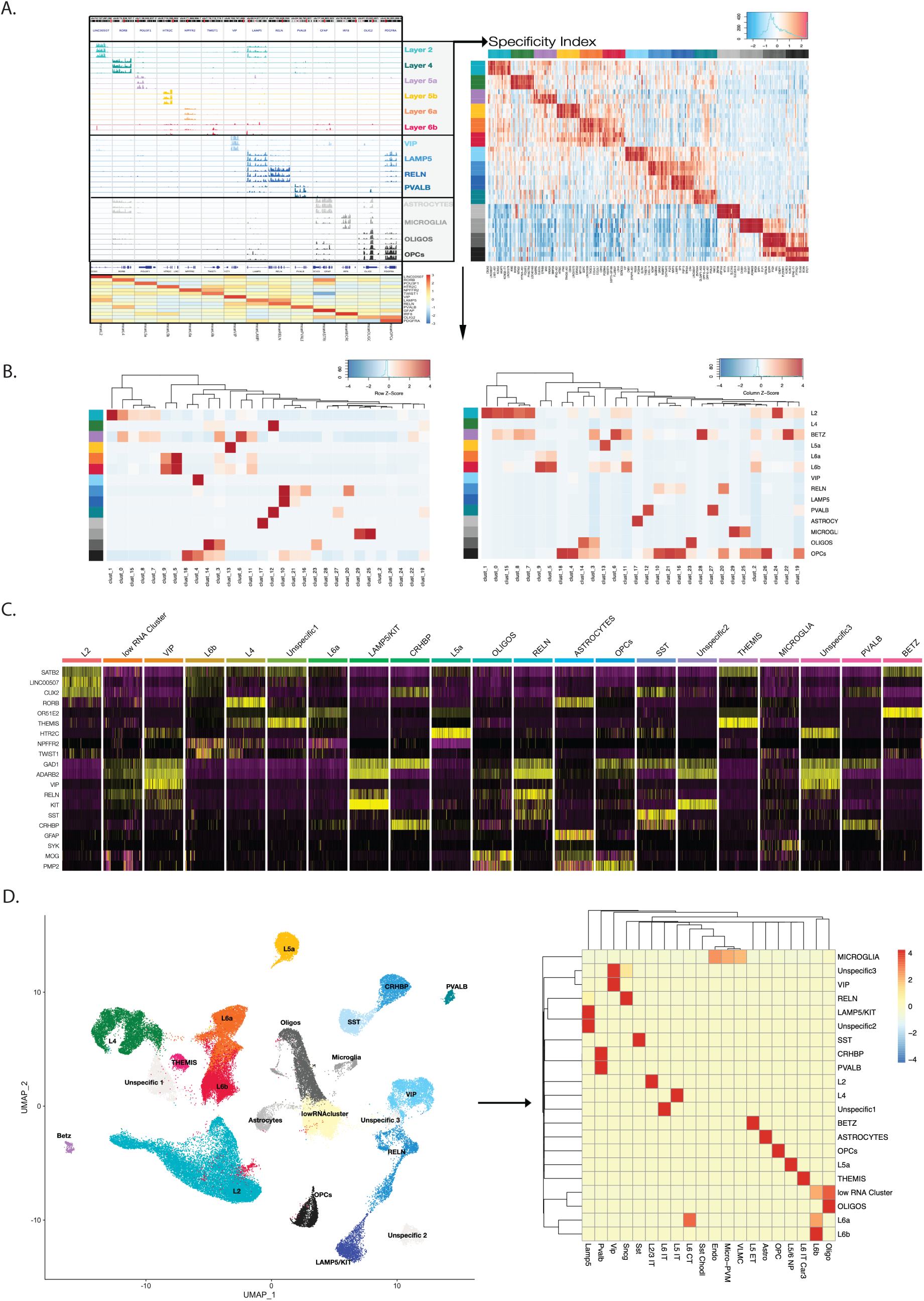
Schematic representation of the steps for sFANS-guided snRNAseq cluster assignment (**A**) Utilization of sFANS derived RNAseq data for the calculation of the cell-type specific Specificity Index for all genes. The heatmap on the right shows the top 20 genes rank ordered through the specificity index analyses for all sFANS isolated cell-types. (**B**) Generation of correlation heatmaps between snRNAseq clusters using the cell-type specific Specificity Index gene ranking, heatmaps scaled to rows and scaled to columns are shown side by side. (**C**) Known marker gene expression within each cluster, as shown on the heatmap, utilized for refinement and validation of cluster assignment. (**D**) UMAP with cluster assignments and color coding based on sFANS-guided cell-type designations and marker gene expression (left panel). Scaled to row correlation heatmap shows correlation between sFANS-guided clusters and clusters as defined in the publicly available human motor cortex datasets (Bakken et al., 2021) (right panel).

**Supplementary Figure 4.**
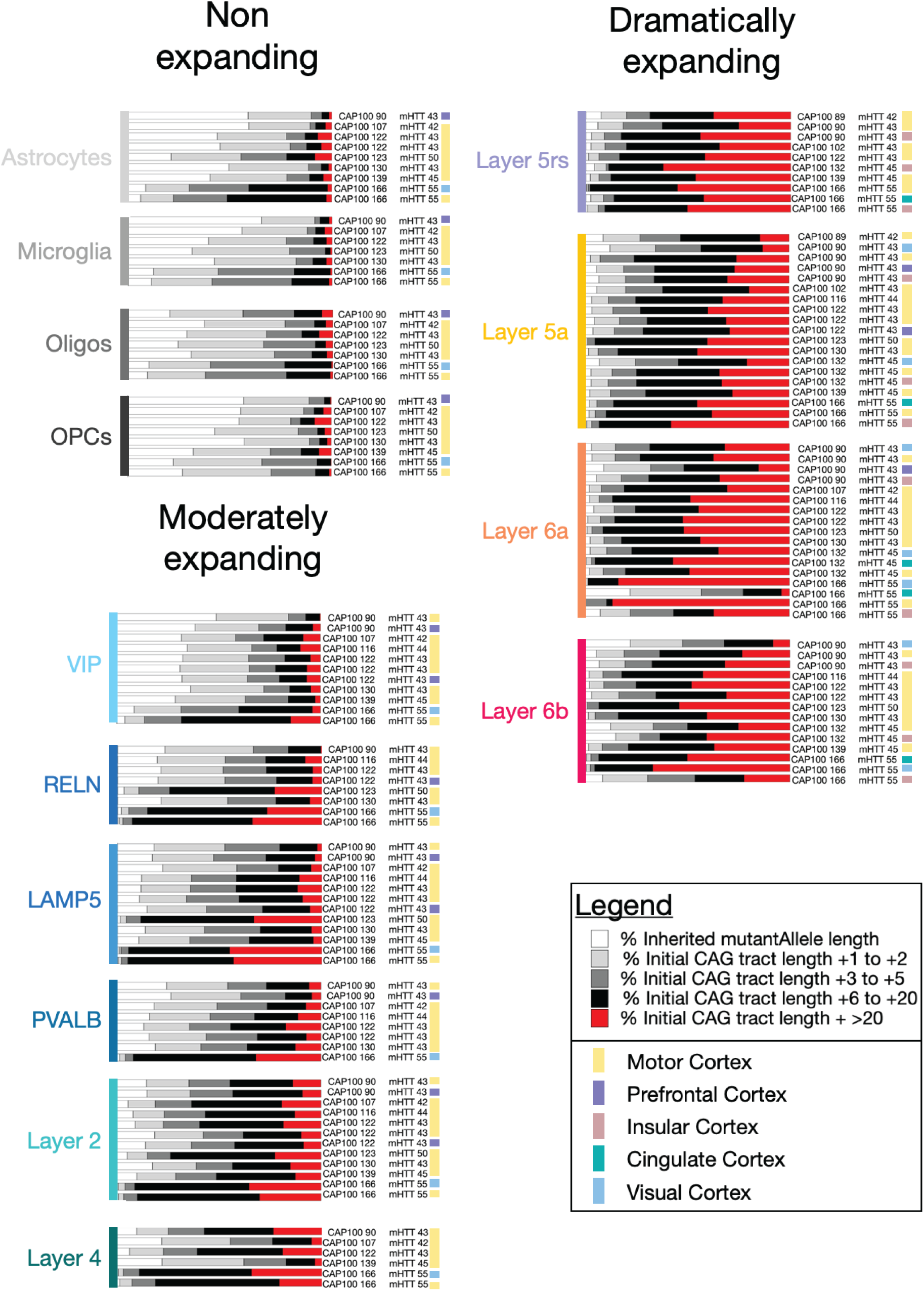
Histograms show the percentage of fragment lengths, detected through analyses of CAG expansion assay data. Data from multiple regions and multiple HD donors is shown for each cell-type. Each horizontal tract represents one cell-type specific sample, to the right of each tract, each sample donor’s inherited mHTT length as well as region of sample origin are indicated. The percentage of fragments detected at the modal mutant allele length is shown in white, the percentage of nuclei where the *mHTT* CAG tract has expanded was calculated and is visualized as the percentage of expansion beyond the modal length by 1 to 2 repeats (light grey), 3 to 5 repeats (grey), 6 to 20 repeats (black), or by more than 20 repeat units (red).

**Supplementary Figure 5.**
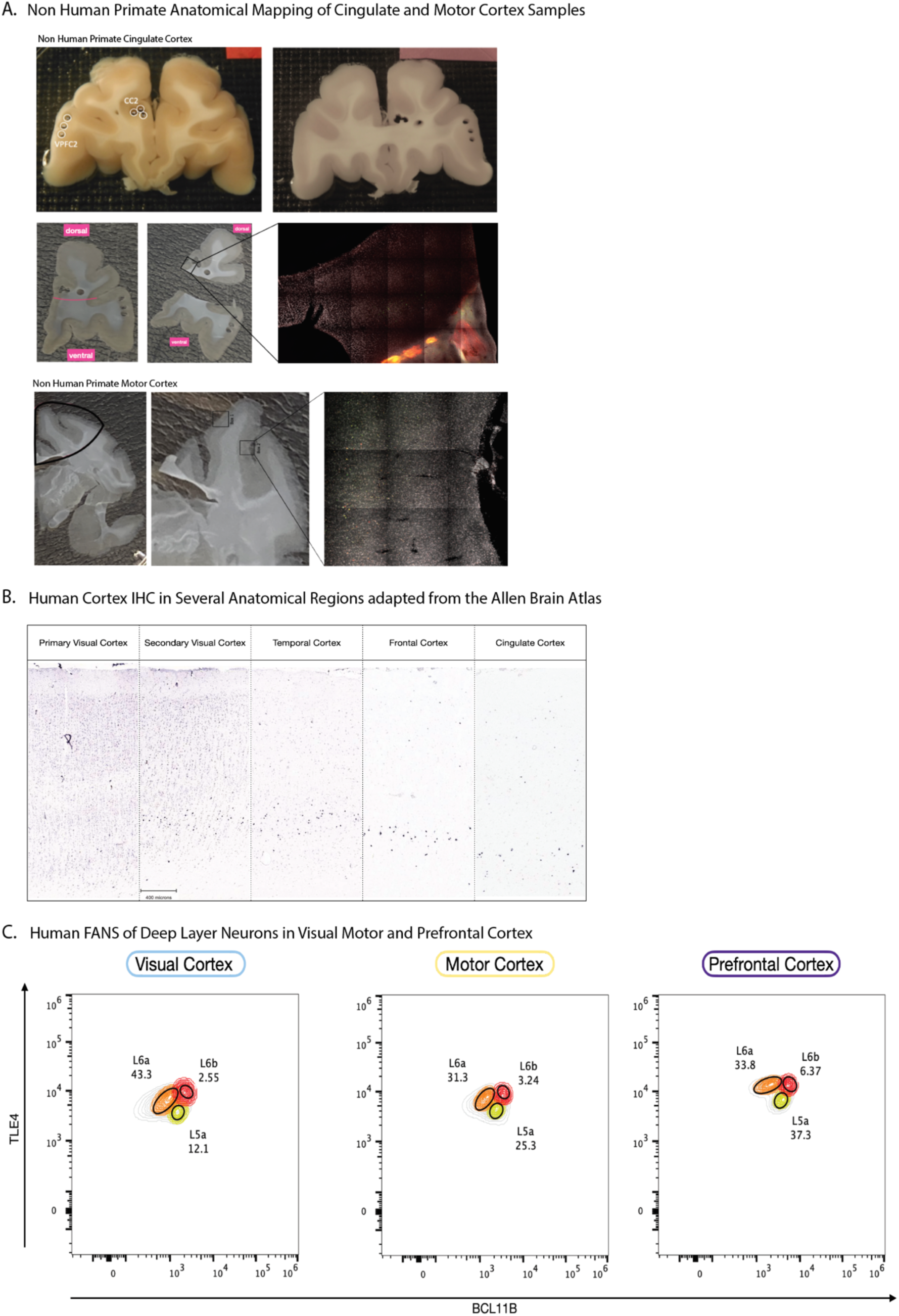
(**A**) Anatomical Mapping of experimental locations in non-human primate ISH experiments. Shown are examples for cingulate and motor cortex containing sections. (**B**) *In situ* hybridization (ISH) data showing expression of *HTR2C* across multiple regions of human cortex, including primary and secondary visual, temporal, frontal, and cingulate cortex. (**C**) sFANS plots showing the distribution of SATB2 high nuclei along TLE4 and BCL11B and relative abundance of layer 5a, layer 6a, and layer 6b nuclei in visual, motor, and prefrontal cortex.

**Supplementary Figure 6.**
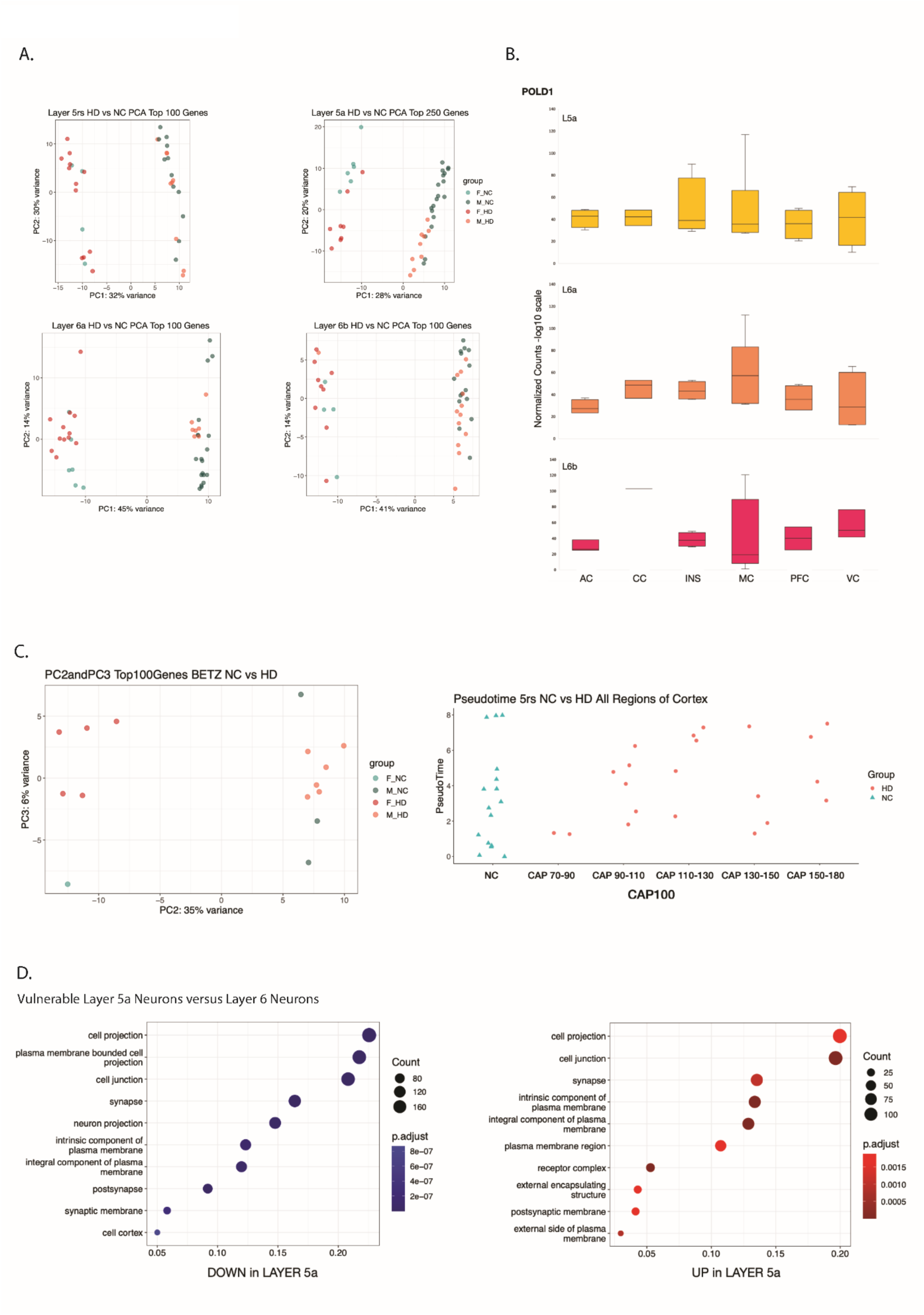
(**A**) Principal Component analyses results for samples from HD and control donors are shown for layer 5rs using the top 100 most variable genes, layer 5a using the top 250 most variable genes, layer 6a using the top 100 most variable genes, and layer 6b using the top 100 most variable genes. PCA plots are color coded by gender and group status (female control (F_NC), male control (M_NC), female HD (F_HD), male HD (M_HD). (**B**) Normalized counts for POLD1 shown on -log10 scale for layer 5a, layer 6a, and layer 6b samples across six cortical regions. (**C**) Principal component analyses of the motor cortex specific Betz cell samples from HD and control donors showing principal components two and three using the top 100 most variable genes (left). Bulk pseudotime analyses results for layer 5rs depicting molecular responses along the pseudotime axis in control samples and in HD samples across CAP scores. (**D**) Results from Gene Ontology Cellular Component pathway enrichment analysis of genes downregulated or upregulated in HD layer 5a versus layer 6 populations (p adj < 0.05 by DESeq2 and filtered for genes with accessible promotors detected by ATACseq).

**Supplementary Figure 7.**
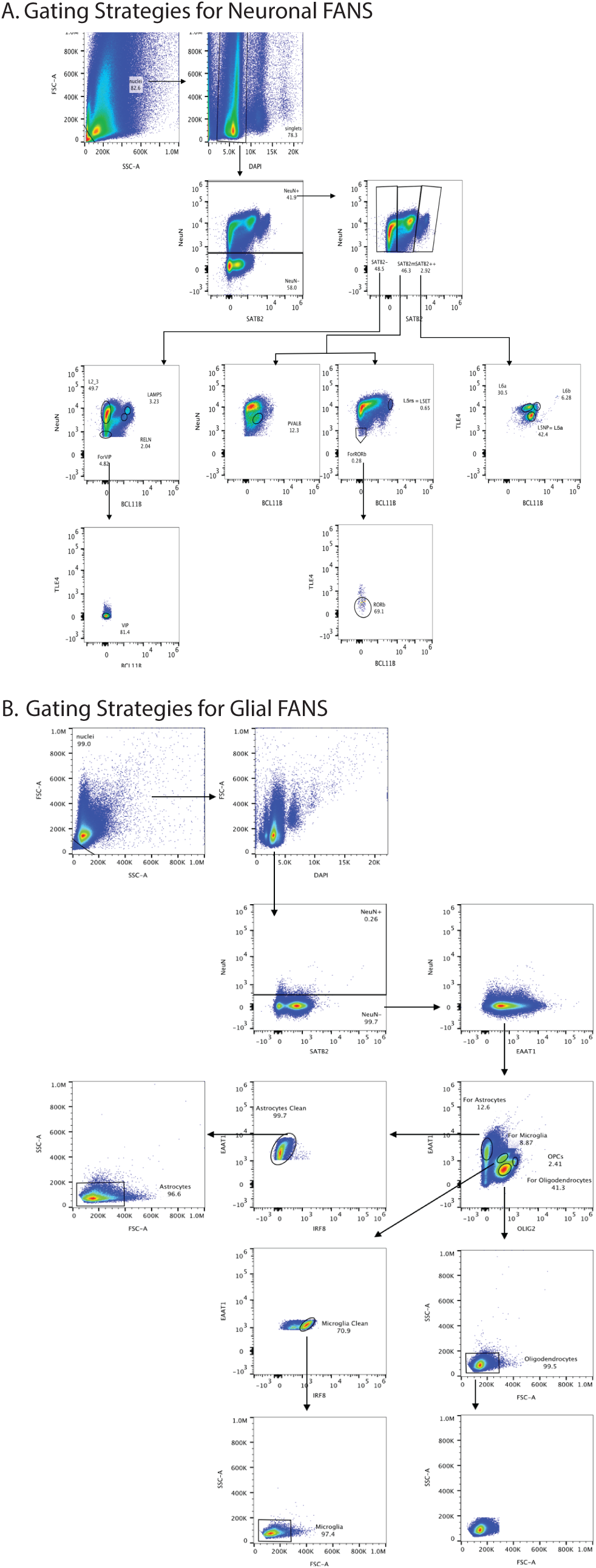
(**A**) Gating Strategies for Neuronal sFANS. Sequential steps for the isolation of neuronal cell-types are shown by the arrows. (**B**) Gating Strategies for Glial sFANS. Sequential steps for the isolation of glial cell-types are shown by the arrows.

## STAR METHODS

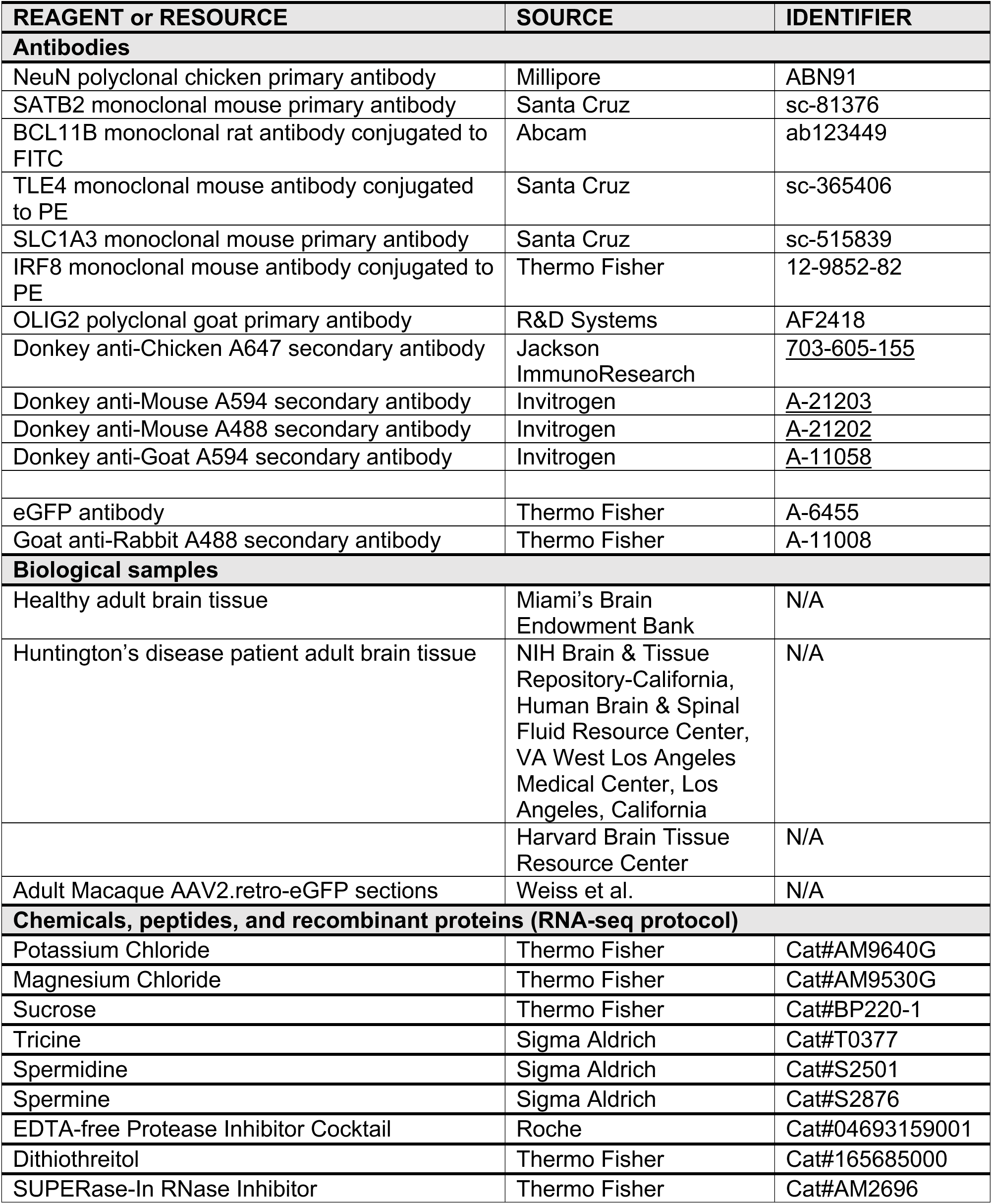

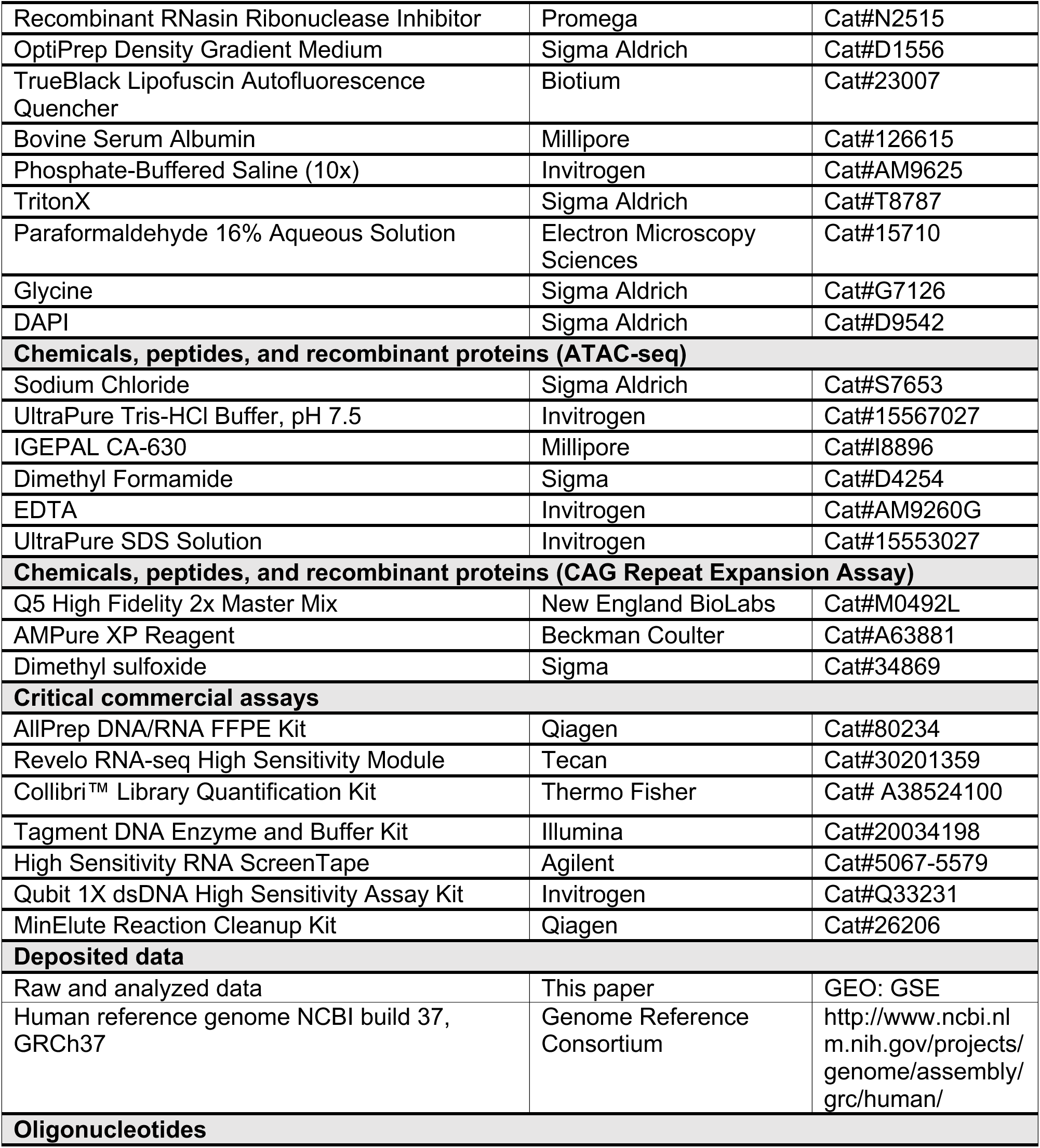

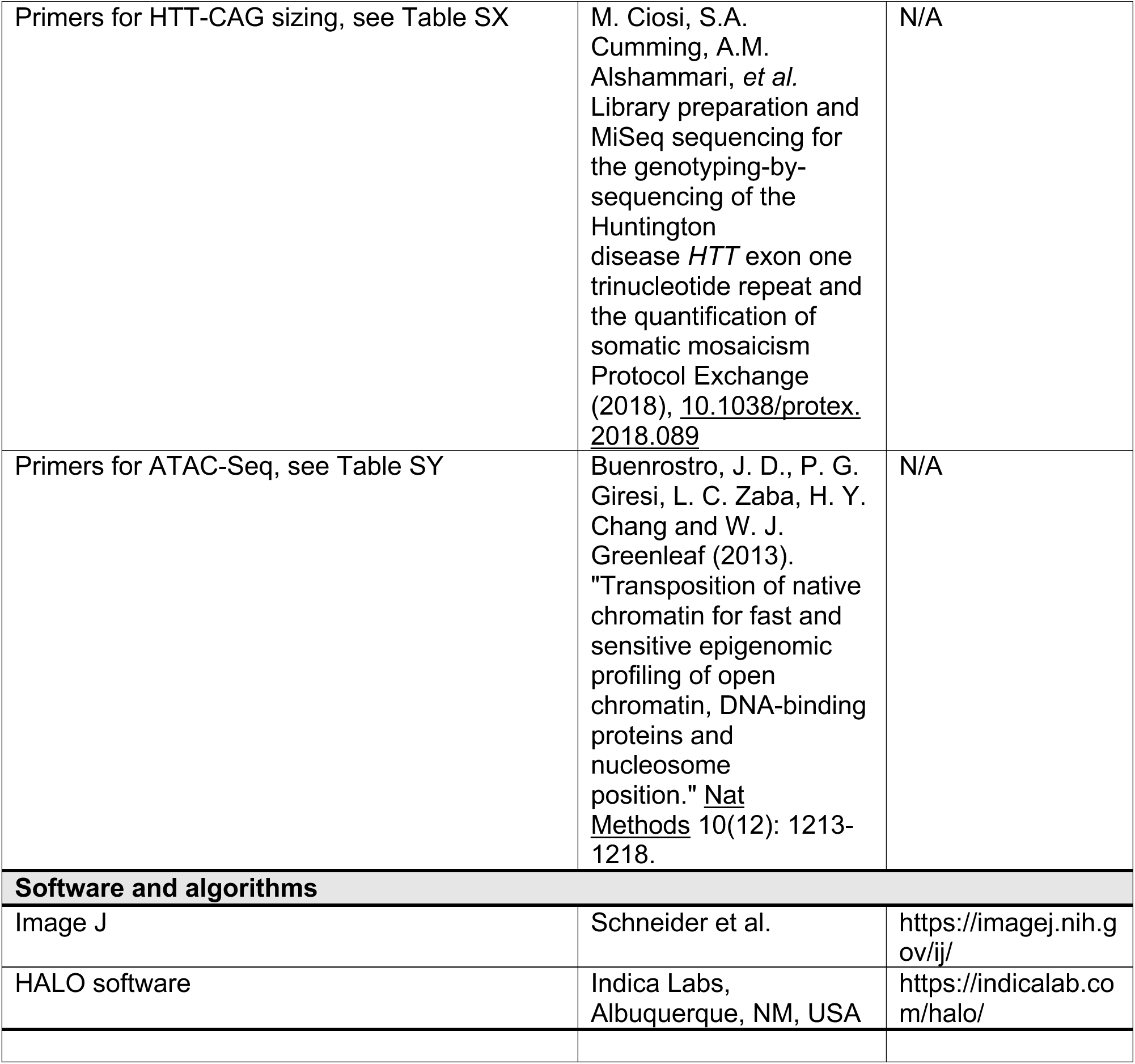
Key resources table.

## MATERIALS AND METHODS

### Sample Collection

De-identified human post-mortem brain tissue analyzed in this study was determined to be exempt from Institutional Review Board (IRB) review according to 45 CFR 46.102 (f). For this work, brain samples were obtained from Miami’s Brain Endowment Bank™ or through the NIH NeuroBioBank and sourced from either the Harvard Brain Tissue Resource Center (HBTRC) or the NIH Brain & Tissue Repository-California, Human Brain & Spinal Fluid Resource Center, VA West Los Angeles Medical Center, Los Angeles, California (Rothenberg et al.), which is supported in part by National Institutes of Health and the US Department of Veterans Affairs. Inclusion criteria were; a postmortem interval (Hernandez-Garcia et al.) of less than 24 hours, a RIN number of greater than 6, male or female sex, all races and an age range between 20-70 years old. Drug addiction and schizophrenia as well as clinical evidence of brain cancers were reason for sample exclusion. While not preferred, samples from donors with a history of other non-brain cancers and diabetes were accepted and the tissue should not have been thawed and re-frozen. Fresh frozen (FF) as well as formalin-fixed and paraffin embedded (FFPE) specimen were obtained from multiple donors and several Brodmann areas. Cortical samples used in this work were derived from primary motor cortex (MC, Brodmann area BA4), prefrontal cortex (PC, Brodmann area BA9), cingulate cortex (CC, Brodmann area BA24), insular cortex, auditory cortex, specifically Wernicke’s area (AC, Brodmann area BA22), visual cortex (VC, including Brodmann areas BA17, as well as Brodmann areas 18/19). Inclusion criteria for HD brain specimen were as described in the section above with the addition that HD samples from individuals who carried around 40 or more CAG repeats in the first exon of the *HTT* gene were selected and obtained from the brain banks. Vonsattel grades, when available, were defined by the brain banks, following classically assessment of striatal pathology (Vonsattel et al., 1985).

Available metadata for each donor sample used in this study is provided in our metadata tables for tissue donors (Table S1) and include information on brain bank source, sex, age, race, post mortem interval (Hernandez-Garcia et al.), year of death, clinical diagnosis, group Normal Control (NC), Huntington’s Disease (HD), Vonsattel grade (when available), brain weight, and RNA Integrity Number (RIN) (when available), as well as CAG repeat tract length of the normal allele, CAG repeat tract length of the mutant allele, CAG repeat structure of the normal allele, CAG repeat structure of the mutant allele for all HD donors and a select set of NC donors that were genotyped.

### Sample Preparation

Fresh frozen as well as FFPE samples from the requested Brodmann areas were provided by the brain banks and shipped to The Rockefeller University overnight. FFPE sample blocks were shipped at room temperature (RT). FF samples were shipped on dry ice. Upon arrival at The Rockefeller University all samples were photographed and catalogued. FFPE samples were stored at RT and FF specimens were stored at -80°C. For nuclei isolation experiments, FF sample blocks were further sub-dissected. On the day of sub-dissection, sample blocks were removed from minus 80°C and moved to minus 40°C for two hours. After two hours, samples were removed from minus 40°C and placed on a metal plate on dry ice. Sub-dissections were performed on frozen specimen with the goal of i) obtaining samples which contained all layers of cortex and ii) to arrive at an aliquot size of samples weighing between 150 mg and 350 mg. At no point was the tissue allowed to thaw.

### Isolation of Cortical Cell-Types

Procedures for nuclei isolation and sFANS were based on protocols previously established in our lab (Kriaucionis and Heintz, 2009) (Mellen et al., 2012) (Xu et al., 2018) and were applied with modifications. Original protocols had previously been optimized in human samples for the isolation of cerebellar cell-types and have been presented in detail elsewhere (Xu et al., 2018). General procedures for sFANS and for the application of our sFANSeq strategy will be discussed in brief in the following sections, while modifications of the original protocols for the isolation of human neocortical cell-types will be presented in detail.

### Antibodies and Reagents

Seven primary antibodies were utilized to build our multiplexed sFANSeq panel. This multiplexed panel was used for the isolation of all major human cortical cell-types. The seven target proteins and antibodies for the sFANSeq panel included Neuronal Nuclei Antigen (NeuN, Millipore Cat# ABN91, RRID: AB_11205760, used at a concentration of 1:500), Special AT-Rich Sequence Binding Protein 2 (SATB2, Santa Cruz Biotechnology Cat# sc-81376, RRID:AB_1129287, used at a concentration of 1:100), BAF Chromatin Remodeling Complex Subunit (BCL11B aka CTIP2, Abcam Cat# ab123449, RRID:AB_10973033 conjugated to FITC, used at a concentration of 1:100), Transducin Like Enhancer Of Split 4 (TLE4, Santa Cruz Biotechnology Cat# sc-365406, RRID:AB_10841582 conjugated to PE, used at a concentration of 1:100), Solute Carrier Family 1 Member 3 (SLC1A3, Santa Cruz sc-515839 used at a concentration of 1:2000), Interferon Regulatory Factor 8 (IRF8, Thermo Fisher Scientific Cat# 12-9852-82, RRID:AB_2572742 conjugated to PE, used at a concentration of 1:100), Oligodendrocyte Transcription Factor 2 (OLIG2, R and D Systems Cat# AF2418, RRID:AB_2157554, used at a concentration of 1:500). Combination of these seven antibodies into a multiplexed multi-color panel, the use of hierarchical and Boolean logic and considering antibody labeling strength, as well as serial sorting and serial staining strategies allowed for the isolation of up to 16 distinct cell-types from one human cortical sample aliquot. Detailed information on the specific primary and secondary antibodies used in this work is provided in the key resources table.

### Nuclei Isolation

Sample aliquots were thawed on ice for approximately 20 minutes. Samples were transferred into homogenization buffer and homogenized through dounce-homogenization and application of 30 strokes with the loose pestle A, followed by 30 strokes with the tight pestle B as described by Xu et al. (Xu et al., 2018). The homogenate was brought to 24% iodixanol concentration and layered on top of a 27% iodixanol cushion for density gradient centrifugation. Centrifugation was performed for 30 minutes in an SW41 rotor in a Coulter Beckman ultracentrifuge at 4°C, 10,000 RPM, with maximum acceleration and slowest deceleration. Pelleted nuclei were harvested from the bottom of the ultracentrifuge tube and resuspended in homogenization buffer.

### Nuclei Preparation for Fluorescence Activated Nuclei Sorting

Except for those samples intended for snRNAseq, nuclei were fixed by adding 1mL of Homogenization buffer with 1mM EDTA and 1% formaldehyde (PFA) for 8 minutes at room temperature. The reaction was quenched through the addition of 0.125 M glycine. After 5 minutes of incubation, nuclei were washed once in homogenization buffer, followed by a transfer wash into blocking buffer containing Triton 0.05% X-100. Nuclei were blocked and permeabilized in Triton X-100 containing blocking buffer for 30 minutes at RT.

### True Black Treatment

After blocking, nuclei were treated with True Black (TB) to minimize autofluorescence signal. Nuclei were habituated to non-Triton X-100 containing wash buffer to avoid TB wash-out by Triton X-100 and retain suppression of autofluorescence signal through TB. Nuclei were washed twice in Wash buffer without TritonX-100 and resuspended in 100 µl of 40% ethanol containing TrueBlack Lipofuscin Autofluorescence Quencher (Biotium, #23007) for 40-50 seconds.

### Antibody Staining of Nuclei

After isolation, nuclei were resuspended in resuspension buffer in 100μl total volume and incubated with primary antibodies listed in the neuronal sFANS panel for 60 minutes at RT. Antibody NeuN was added at a 1:500 concentration, anti-SATB2, anti-BCL11B conjugated to FITC, and anti-TLE4 conjugated to PE were added at a concentration of 1:100. After 60 minutes, nuclei were washed three times by pelleting (3min, 1000 x g) and resuspension in wash buffer. Secondary antibodies were added to the nuclei in 500μl total volume and at a dilution of 1:1000, for a 30-minute incubation at RT. Detailed information on the specific primary and secondary antibodies used in this work is provided in the antibody staining strategy table. Anti-NeuN antibodies were labeled with anti-chicken 2° antibodies conjugated to a A647 fluorophore. Anti- SATB2 antibodies were labeled with anti-mouse 2° antibodies conjugated to a A594 fluorophore. The concentration of secondary antibodies was kept at 1:1000 in a total volume of 500μ wash buffer and nuclei were incubated for 30 minutes. Three washes in wash buffer were followed by the last resuspension of nuclei in a DAPI (1:25,000) containing wash buffer solution. Nuclei were separated by FACS immediately or stored overnight (ON) at 4°C.

**Table.** Antibody Staining Strategy.

| ANTIBODIES |  |  |  |  |  |  |  |  |  |
| --- | --- | --- | --- | --- | --- | --- | --- | --- | --- |
|  | Probe Name | Vendor | Catalog Number | Type | Clonality | Host Species | Target Species | Gene | Fluorophore |
| ALL FANS | NeuN Ab | Millipore | ABN91 | primary antibody | Polyclonal | Chicken | / | RBFOX3 | / |
|  | Donkey anti-Chicken A647 |  |  | secondary antibody |  | Donkey | Chicken | / | A647 |
| Neuronal FANS | SATB2 Ab | Santa Cruz | sc-81376 | primary antibody | Monoclonal | Mouse | / | SATB2 | / |
|  | Donkey anti-Mouse A594 |  |  | secondary antibody |  | Donkey | Mouse | / | A594 |
|  | BCL11B Ab | Abcam | ab123449 - FITC | conjugated antibody | Monoclonal | Rat | / | BCL11B | FITC |
|  | TLE4 Ab | Santa Cruz | sc-365406 PE | conjugated antibody | Monoclonal | Mouse | / | TLE4 | PE |
| Glial FANS | SLC1A3 Ab | Santa Cruz | sc-515839 | primary antibody | Monoclonal | Mouse | SLC1A3 | SLC1A3 |  |
|  | Donkey anti-Mouse A488 |  |  | secondary antibody |  |  |  |  |  |
|  | IRF8 Ab | Thermo Fisher | 12-9852-82 PE | conjugated antibody | Monoclonal | Mouse | IRF8 | IRF8 | PE |
|  | OLIG2 Ab | R&D Systems | AF2418 | primary antibody | Polyclonal | Goat | OLIG2 | OLIG2 |  |
|  | Donkey anti-Goat A594 |  |  | secondary antibody |  |  |  |  |  |

### Flow Cytometry – Serial Sorting, Post-sorting, and Staining Strategies

Boolean logic and serial sorting strategies were applied to maximize the number of cell-types obtainable from each sample aliquot. Sorting strategies were developed to achieve highest efficiency within the inherent constraints of possible fluorophore combinations and technical boundaries of The SONY MA900 Fluorescence Activated Cell Sorting (FACS) instrument. The SONY MA900 was utilized for all experiments described in this work. The first sort, henceforth referred to as the separation sort, was performed to separate non-neuronal nuclei (NeuN-), from three groups of mixed neuronal nuclei populations (NeuN+SATB2-, NeuN+SATB2medium, NeuN+SATB2+). The separation sort was followed by 1) post-separation staining procedures on NeuN negative nuclei and 2) sub-sorts of the three previously sorted NeuN positive nuclei containing tubes. NeuN negative nuclei underwent post-staining procedures, including glia-specific primary and secondary antibody incubations as well as wash steps that were equivalent to the steps described above. Detailed instructions and gating steps for our sFANSeq strategy will be provided below.

### sFANS Gating Strategy, Data Processing, Visualization, and Quantification sFANS Gating and Sorting Strategies

sFANS data pre-processing, analyses, and visualization were performed using the SONY visualization tool as well as the FlowJo™ v10.8 Software (BD Life Sciences) (FlowJo™ Software for Mac. Ashland, OR: Becton, Dickinson and Company; 2023). sFANS .fcs files were loaded into FlowJo, and gates were drawn. Fluorophore measurements were plotted in biexponential mode.

The multi-step gating strategy to isolate ten neuronal cortical cell-types included:

1. Size selection of nuclei was performed by excluding particles below approximately 100K along the forward scatter area (FSC-A) and side scatter area (SSC-A) axes and smaller than the majority of other particles measured.
2. Doublet discrimination and singlet selection was performed by gating on the lowest positive population along the liner DAPI axis.
3. Separation of NeuN positive neuronal nuclei from NeuN negative non-neuronal nuclei was performed along the NeuN axis.
4. NeuN positive nuclei were plotted along the NeuN and SATB2 axes and divided into three populations NeuN positive populations:

A. NeuN+/SATB2- nuclei, a population located below the 10^2^ mark along the SATB2 axis. This group of nuclei was plotted against NeuN and BCL11B, revealing four subpopulations.

1. NeuN+low/SATB2-/BCL11B- nuclei are enriched for VIP expressing interneurons. This VIP expressing population was further purified by plotting and gating against nuclei that spread upwards along the TLE4 axis.
2. NeuN+high/SATB2-/BCL11B- nuclei are enriched for CUX2 expressing cortical layer 2 excitatory neurons.
3. NeuN+high/SATB2-/BCL11Bmedium nuclei are enriched for a RELN expressing inhibitory neuron population.
4. NeuN+high/SATB2-/BCL11B+ nuclei are enriched for a LAMP5 expressing inhibitory neuron population.
5. B. NeuN+/SATB2medium nuclei, a population located above the 10^2^ mark along the SATB2 axis and separated below the highest SATB2 positive population. NeuN positive SATB2 medium nuclei were further divided into NeuN+/SATB2mediumlow and NeuN+/SATB2mediumhigh populations and plotted against NeuN and BCL11B.

1. NeuN+low/SATB2mediumlow/BCL11B+ nuclei are enriched for PVALB expressing inhibitory neurons.
2. NeuN+low/SATB2mediumhigh/BCL11B- nuclei are enriched for layer 4 excitatory neurons. This RORb expressing population was further purified by plotting and gating against nuclei that spread upwards along the TLE4 axis.
3. NeuN+high/SATB2mediumhigh/BCL11B+high is a small region-specific population, enriched for the Betz or Von Economo neuron transcription factor POU3F1. Populations isolated from this location are region-specific and are henceforth referred to as layer 5rs to allow for consistent comparisons across all regions.
4. A second population within the SATB2 medium high range appeared NeuN+low/SATB2mediumhigh/BCL11B+high and was only inconsistently present across samples from the same regions. Due to its inconsistency, it has been difficult to define this population. However, since this population may represent a rare and potentially region-specific CRHBP expressing interneuron population it will continue to be of interest to further dissolve and define this population.

1. C. NeuN+/SATB2+, a population forming at the highest end along the SATB2 axis, contained three populations within it.

1. NeuN+/SATB2+/BCL11B+/TLE4+ nuclei were enriched for *HTR2C* expressing layer 5a excitatory neurons.
2. NeuN+/SATB2+/BCL11B+/TLE4+high nuclei were enriched for CTGF and NPFFR2 expressing layer 6a excitatory neurons.
3. NeuN+/SATB2+/BCL11B+high/TLE4+high nuclei were enriched for CTGF expressing layer 6b excitatory neurons. This cell population is sparse, does not express NPFFR2, and specific marker genes appear variable across the different regions of cortex.

Gating strategy to isolate four non-neuronal cell-types:

1. Previously separated NeuN negative nuclei underwent size selection for nuclei using the FSC and SSC scatter, as described above.
2. Single nuclei selection was performed through assessment of DAPI staining intensity as described.
3. NeuN negative nuclei staining intensity was plotted for the non-neuronal antibody markers utilized and with the goal to isolate four NeuN negative populations:

A. NeuN negative EAAT1 (aka SLC1A3) positive nuclei, for astrocytes
B. NeuN negative IRF8 positive nuclei, for microglia
C. NeuN negative OLIG2 positive nuclei, for oligodendrocytes
D. NeuN negative OLIG2 high positive nuclei, for OPCs

Supplementary Figure7 shows our gating strategy for the isolation of neuronal nuclei and non-neuronal nuclei from one control donor motor cortex sample.

### sFANS Data Processing, Visualization, and Quantification Methods

sFANS data were processed using the FlowJo™ v10.8 Software (BD Life Sciences) (FlowJo™ Software for Mac. Ashland, OR: Becton, Dickinson and Company; 2023). For population quantifications, data were batch processed followed by individual sample inspection and gate correction as warranted. Gating plots were produced in batches using the layout editor function. Raw data for quantification analyses were processed in batches using the table editor function and results were extracted in .csv format. Quantification data was read into R-Studio for statistical analyses and to produce sFANS population quantification plots.

### DNA and RNA Extraction, Purification, and cDNA Library Construction from Nuclear RNA

For RNAseq experiments, RNA was extracted from fixed and sorted nuclei using the Qiagen AllPrep DNA/RNA FFPE kit (Qiagen, #80234). The original Qiagen RNeasy FFPE kit protocol had been previously optimized for our sample type. DNA from all samples was isolated and purified Genomic DNA was purified using AllPrep DNA/RNA FFPE Kit (Qiagen, #80234), purified DNA was stored at -20°C.

### RNA Sequencing

Libraries were assessed for concentration and quality using the Qubit and Tapestation assays. Samples were pooled and sequenced using the NovaSeq SP 2x150 with the goal to reach a read depth of approximately 40 million reads per sample.

### Nuclei Preparation for Single Nucleus RNA Sequencing

Nuclei intended for 10X snRNAseq experiments remained unfixed but were otherwise treated as described above and identically to nuclei intended for all other down-stream applications. Further processing of nuclei isolated for snRNAseq experiments was performed by The Rockefeller University’s Genomics Resource Center. In brief, the number of nuclei per sample was counted and the appropriate loading amount was calculated. Nuclei were loaded and processed on the 10X platform for snRNAseq. 1000121 Chromium Next GEM Single Cell 3’ GEM, Library & Gel Bead Kit v3.1, 16 rxns, 1000215 Dual Index Kit TT Set A 96 rxns, and Chromium Single Cell Controller were used. Before deep sequencing, the resulting libraries were pooled in groups of 3- 4 samples and were sequenced on MiSeq using the Nano 300 kit. The MiSeq results provided 1 million reads over all samples and were utilized to assess the ratio of empty droplets and estimate nuclei yield. Finally, all samples were pooled and sequenced on NovaSeq using the S4 flow cell 2x100 kit to generate 8 billion reads over all samples and to allow for sufficient read depth given the total number of nuclei.

### ATACseq of sFANS-Enriched Deep Layer Cortical Neurons

The ATACseq protocol utilized in this work was adapted from Corces et al. (Corces et al., 2017). Sample processing steps will be outlined in brief below. For our ATACseq experiments we focused on four deep layer cell-types of interest and performed nuclei isolations as described above from motor cortex samples of control donors. Following nuclei sorting, nuclei were spun for 5 minutes at 950 g, supernatant was removed and for samples containing up to 50,000 nuclei, 50μl lysis buffer (10 mM Tris-HCl pH 7.6, 10 mM NaCl, 3 mM MgCl2, 0.01% NP-40) was used for resuspension. For samples containing more than 50,000 nuclei, the additional amount of 10μl lysis buffer for every additional 10,000 nuclei was used. Nuclei were spun at 500 g for 10 minutes at 4°C, supernatant was removed, and nuclei were resuspended in 200μl transposition mix (1x TD buffer containing X µl of Illumina Tagment DNA TDE1 Enzyme per every X nuclei). Samples were incubated on a heat block set to 37°C for 30 minutes. After 30 minutes, 150μl reverse X-link solution was added, and nuclei were incubated at 55°C overnight at 500 RPM. On day two, DNA was isolated using the Qiagen MinElute Reaction Cleanup Kit. Next, PCR amplification was performed. PCR amplification reagents were mixed in a clean strip tube before the sample was added. Amplification was performed using the following settings: (72C – 5min, 98C – 30sec, appropriate number of amplification cycles x (98C - 10sec, 63C – 30sec, 72C – 1 min) with NEBNext® High-Fidelity 2X PCR Master Mix (New England Biolabs, #M0541S) and barcoded Nextera primers (1.25 μM each) (Buenrostro et al., 2013). The number of amplification cycles was dependent on the number of nuclei contained within the sample. Less than 15,000 nuclei, 15 cycles were used, for samples containing between 15,000 and 100,000 nuclei, 13 cycles were used. The maximum input was kept below 75,000 nuclei for 13 PCR cycles. Upon completion of the library amplification steps, double sided bead purification was performed. Qubit and Tapestation were used to confirm correct library sizing and sufficient sample concentrations before pooling. Samples were pooled for a minimum concentration of 2.6 nM total in 100μl volume and ATACseq libraries were sequenced using NovaSeq6000 SP 2x100.

### Sample Sectioning for Immunohistochemistry and *In-situ* Hybridization

FFPE samples were sectioned on a Leica HistoCore AUTOCUT Rotary Microtome. Fresh frozen tissue was sectioned on a Leica CM3050 S Cryostat. Samples were mounted on SuperFrost™ Plus and FFPE slides were stored at RT, whereas fresh frozen sections were stored at -20°C. Nonhuman primate (NHP) samples were provided by our collaborator Dr. Jodi McBride. Details on tissue processing as performed by our collaborators as well as IHC methods for GFP labeling were as described by Weiss et al. (Weiss 2020). Photo documentation, as provided by our collaborator and anatomical landmarks (Supplementary Figure S5A) were used to define the anatomical regions contained within the sections used in our experiments.

### Immunohistochemistry

Standard operating procedures were based on IHC protocols sourced from the BioLegend website and applied with modifications. In brief, FFPE slides were deparaffinized with xylene, dehydrated in baths of decreasing EtOH concentrations. Endogenous peroxidase activity was blocked through incubation with 3% 2 in methanol and slides were rinsed. Antigen retrieval was performed in citrate buffer (pH 6.0) heated to 95-100°C for 10 minutes. Slides were set aside for cooling for 20 minutes and rinsed two times with PBS. Next, slides were incubated in blocking buffer containing 10% normal donkey serum in PBS with 4 drops Avidin per ml buffer. Sections were coverslipped and incubated in a humidified chamber at RT for one hour. Blocking buffer was drained from the slide and primary antibodies in the appropriate dilutions (1:100) in dilution buffer containing 1% normal donkey serum in PBS with 4 drops Avidin per ml buffer. Sections were coverslipped and incubated in a humidified chamber at 4°C overnight. On day two, slides were washed with PBS followed by incubation of biotinylated secondary antibody (1:200) in dilution buffer. Sections were coverslipped and incubated in a humidified chamber at RT for 30 minutes. Slides were washed with PBS and Streptavidin-HRP conjugate (NEL-700 NEN Life Science Products) (1:100) in 1% normal donkey serum containing PBS was applied. Sections were coverslipped and incubated in a humidified chamber at RT for 30 minutes and protected from light. Slides were washed and incubated with TSA (Tyramide Signal Amplification Biotin System, NEL-700 NEN Life Science Products) diluted to a 1:100 concentration in amplification diluent. Sections were coverslipped and incubated in a humidified chamber at RT for 10 minutes. Slides were washed with PBS and finally, slides were cover-slipped and incubated at RT with ABC (ABC Kit Elite PK-6100, Standard Vector Labs) for 30 min at RT. Slides were washed and DAB substrate solution (0.05% DAB, 0.015% 2 in PBS, 2 DAB kit tablets, 1 gold & 1 silver in 15ml d) was applied to the slides, using a disposable pipet. Development time was generally kept below 5 min and sustained until the desired color intensity was reached. Slides were washed with PBS, mounted with aquamount, and dried overnight before imaging was performed.

IHC experiments for visualization of AAV2.retro treated NHP sections were performed following the methods described in Weiss et al. (Weiss et al., 2020). In brief, the 40 micron thick sections were incubated with an antibody against eGFP (Thermo Fisher, A-6455, 1:1000) over night at room temperature followed by one hour incubation with a goat anti rabbit secondary Alexa Fluor A488 (Thermo Fisher, A-11008, 1:500) conjugated secondary antibody.

### *In-situ* Hybridization

*In-situ* hybridization experiments performed over the course of this work were executed using the commercially available RNAscope® assay kit. Multiplexed fluorescent ISH were performed following the vendor’s protocols according to the tissue type used. Tissue types used in this work were limited to fresh frozen NHP sections. The following ACD protocols were used: User Manual RNAscope® 2.5 Multiplex Fluorescent Reagent Kit v2 Assay (With Sample Preparation and Pretreatment, Document Number 323100-USM), Sample Preparation Technical Note for Fixed Frozen Tissue Using RNAscope® 2.5 Chromogenic Assay (Singleplex and Duplex) TN- 320534/Rev B). Protocol adaptations for the 40-micron thick fixed frozen NHP sample sections were as follows: Sections were mounted onto Superfrost Plus slides and dried at RT for 1 hour. Slides were dipped in water 3 times and dried at RT for one hour followed by a backing step at 60°C for 1 hour. Post-fixation was performed using 4% PFA at 4°C for 1 hour. Slides were dehydrated with an increasing EtOH gradient, and slides were dried at 60°C for 15 minutes. Next, samples were incubated with RNAscope hydrogen peroxide at RT for 10 minutes or until no active bubbling was observed. Slides were rinsed with water and dried for 15 minutes at 60°C. Target retrieval was performed in 1X TR buffer for 10 minutes at 100°C using a steamer apparatus. Slides were rinsed, briefly dipped in 100% EtOH and dried at 60°C for 10 minutes or longer, until dry. The hydrophobic barrier was drawn on the slide, enclosing the tissue sample and Protease III was applied for digestion for 30 minutes at 40°C. Slides were quickly washed with dH2O before the samples were further processed using the appropriate RNAscope® protocol.

### Imaging and Analyses of IHC and ISH Sections

All imaging of non-fluorescent samples was performed on the NanoZoomer-SQ Digital slide scanner (Hamamatsu C13140-01) and using the Hamamatsu NDPScanMonitor software program. Imaging of fluorescent samples was performed on the LSM700 Zeiss Confocal Microscope, using the Zeiss ZEN microscopy software. IHC imaging results from experiments using our cortical neuronal sFANS panel antibodies on FFPE sections and using the above described IHC DAB protocol are not shown. ISH imaging data was inspected and processed for image overlays using ImageJ (Schneider et al., 2012). ISH images were read into the HALO software (Indica Labs, Albuquerque, NM, USA). Signal for GFP and *HTR2C* probes were automatically evaluated by the software and three categories of colocalization signal were extracted to generate anatomical maps: cells strongly positive for eGFP, cells strongly positive for both eGFP and *HTR2C*, and cells strongly positive for eGFP and weakly positive for *HTR2C*. localization maps were generated. Out of all GFP positive cells, the percentage of double positive cells high in eGFP and in *HTR2C*, as well as the percentage of double positive cells high in eGFP and with any *HTR2C* signal intensity were calculated and plotted in one bar chart showing motor and cingulate cortex sample results.

### RNAseq Data Processing

Sequence and transcript coordinates for human hg38 UCSC genome and gene models were retrieved from the BSgenome.Hsapiens.UCSC.hg38 Bioconductor package (version 1.4.1) and TxDb.Haspiens.UCSC.hg38.knownGene Bioconductor libraries (version 3.4.0) respectively. RNAseq reads were aligned to the genome using Rsubread’s subjunc method (version 1.30.6) (Liao et al., 2013) and exported as bigWigs normalized to reads per million using the rtracklayer package (version 1.40.6). Reads in genes were counted using the featurecounts function (Liao et al., 2014) within the Rsubread package against the full gene bodies for nuclear RNAseq (Patro et al., 2017). Normalization and rlog transformation of raw read counts in genes were performed using DESeq2 (Love et al., 2016) (Love et al., 2018) (version 1.20.0) and between sample variability was assessed with hierarchical clustering and heat maps of between sample distances implemented in the pheatmap R package (1.0.10).

### ATACseq Data Processing

The ATACseq reads were aligned with the hg38 genome from the BSgenome.Hsapiens.UCSC.hg38 Bioconductor package (version 1.4.1) with Rsubread’s align method in paired-end mode. Fragments between 1 and 5,000 base pairs long were considered correctly paired. Normalized, fragment signal bigWigs were created with the rtracklayer package. Peak calls were made with MACS2 software in BAMPE mode (Feng et al., 2012) (Zhang et al., 2008). For each deep layer excitatory neuron cell-type of motor cortex the ATAC-seq consensus peaks were called from ATAC-seq datasets generated from control donor samples of BA4 motor cortex. These datasets included layer 5rs (aka Betz cells in BA4) (n=3), layer 5a (n=4), layer 6a (n=4), and layer 6b (n=4), Differential ATACseq signals were identified with the DESeq2 package (Love et al., 2016) (Love et al., 2018) (version 1.20.0). High confidence consensus peaks were derived by creating a nonredundant peak set for each cell type and disease state, and then filtering down to peaks that were present in the majority of samples. These were then annotated to TSS based on proximity using the ChIPseeker package (version 1.28.3) (Wang et al., 2022). NCBI Refseq hg38 gene annotation was used (version 109.20211119).

RNAseq and ATACseq data were inspected using the Integrative Genomics Viewer (Robinson et al., 2011). RNAseq average raw read counts were processed using the DESeq2 package (Love et al., 2016) (Love et al., 2018) (version 1.36.0). Reads were converted into normalized counts by DESeq2 before differential gene expression analysis. Normalized POLD1 counts on a -log10 scale were plotted using bar graphs for layer 5a, layer 6a, and layer 6b samples separately and across six cortical regions. For deep layer cell-types, differential gene expression analysis were performed for each cell-type separately. PCA plots were generated using the transformed, variance stabilized data using the vst() function and plotted using the plotPCA function. MA plots were generated using the ggplot() function to show log2 foldchange and log2 basemean of differentially expressed genes. Differential RNAseq gene expression analysis results were filtered using our cell-type specific ATACseq datasets to exclude genes that lacked consensus ATAC peaks over their annotated TSS positions in NCBI Refseq hg38 (version 109.20211119). The filtered lists of differentially expressed genes (DEGs) (p adj < 0.05) with accessible TSS regions were used for Gene set enrichment analysis (Subramanian et al., 2005) of over-representation of Gene Ontology Cellular Compartment (GOCC) terms with clusterProfiler package (Yu et al., 2012) (version 4.4.4). The list of all genes with accessible TSS regions was used as the ‘background’ list for comparison (‘universe’). Gene lists were generated separately for layer 5a, layer 5rs, layer 6a, and layer 6b samples from differential gene expression analyses between HD and NC samples (P adj. < 0.05) for the identification of GOCC pathways containing an over-representation of genes that showed disease-associated up- or downregulation in HD. Differential expression between HD and controls for a set of genes of interest was extracted for each cell-type separately and plotted using the pheatmap R package (1.0.10).

### Bulk-pseudotime Analyses of sFANSeq Data

We generated bulk pseudotime trajectories using sFANSeq data from HD and control donor deep layer cell-types, including layer 5rs, layer 5a, layer 6a, and layer 6b. To do this, the gene expression levels of each sample were aggregated into a single matrix. This counting matrix was loaded into Seurat object with the Seurat::CreateSeuratObject() function, generating a Seurat (v 4.0, https://satijalab.org/seurat/) object for each cell-type separately. Next, data were log normalized, variable features were assessed, data was scaled, and PCAs were generated using Seurat. Next, pseudotime trajectories were modeled using slingshot (https://github.com/kstreet13/slingshot). Results were plotted to show the pseudotime trajectory of HD samples along the range of different CAP100 scores and in relation to control samples.

### snRNAseq Data Processing

The snRNAseq raw data was processed using the count function in Cell Ranger (10X Genomics, version 6.0). Reads were aligned to the human genome (hg38) and including the intron regions, genes were mapped and counted in each nucleus. Counting matrices were generated in Cell Ranger and utilized in downstream analyses. Data was read into R and further processed using Seurat algorithms as well as other Bioconductor packages and custom code.

To estimate and remove ambient RNA, the counting matrices were loaded into SoupX (https://github.com/constantAmateur/SoupX). The fraction of ambient RNA contamination was estimated and corrected by using the default settings. Doublets, defined as multiple nuclei in a single droplet, were identified by using Scrublet (https://github.com/swolock/scrublet). Details on rationale and computational steps for removal of ambient RNA can be found here: https://bioconductor.org/books/release/OSCA/droplet-processing.html.

Cell cycle phase was estimated as a quality control measure by using cyclone, a function of scran (https://bioconductor.org/packages/release/bioc/html/scran.html). For each sample, a Seurat (v 4.0, https://satijalab.org/seurat/) object was created by using the corrected matrices. The previously estimated information on doublet scores and cell cycle stages was also added to the Seurat object. After this step, the proportion of mitochondrial genes for each sample was estimated using the Seurat::PercentFeatureSet() function. Nuclei with mitochondrial contents greater than 1% were removed. Distribution of gene counts (detected), transcript count (sum), and mitochondrial content (subsets_Mito_percent) in each cell was calculated and the outliers, with extremely high/low levels in either of the three criteria, were removed. Then, gene expression was normalized by using Seurat::NormalizeData() with logNormalization and scale factor=10000. And lastly, the top 3000 variable features of each object were identified by using Seurat::FindVariableFeatures() function on the Variance Stabilization Transformed (vst) data.

The pre-processed Seurat objects were integrated into one single object by using the canonical correlation analysis (CCA) workflow. In brief, anchors, which constitute shared variable features across all samples, were identified by using the Seurat::FindIntegrationAnchors() function. Then, all datasets were integrated using Seurat::IntegrateData() function. After integration, the assay named “integrated” was set as the default assay and the integrated object was scaled by using the Seurat::ScaleData() function.

To optimize projection and clustering, we programmatically assessed the different cluster-resolution outcomes when different numbers of PCs were used. Principal component analyses (PCA) were performed using Seurat::RunPCA () with up to 40 PCs. The optimized number of PCs was determined by using the elbow plot. Then, the selected number of PCs was applied to produce projection maps using the Seurat::FindNeighbors() and Seurat::RunUMAP functions. After UMAPs were generated, we tested clustering by Seurat::FindClusters() with resolutions ranging from 0.1 to 1.0. Results were plotted using the clustree function (https://cran.r-project.org/web/packages/clustree/vignettes/clustree.html). Then, the optimal resolution with the most clusters identified and the least crosstalk between cluster lineages was applied to our data.

The merged dataset was converted into SingleCellExperiment format and cell type – clustering was performed by using the SingleR package with default settings (https://bioconductor.org/packages/release/bioc/html/SingleR.html). To annotate cell-types, specificity index results, which provide ranked marker cell-type specific gene lists as described in (Dougherty et al., 2010), for all cell-types isolated through the sFANSeq method from three representative control donor samples were calculated and applied to a reference dataset converted in SingleCellExperiment format. Correlation matrixes were calculated and plotted and snRNAseq clusters were assigned to sFANSeq cell-types based on their correlation scores. For cell-type clusters that were not isolated through our sFANSeq panels, known marker genes as well as cross-correlations with published human motor cortex datasets (Hodge et al., 2019) was performed. Lastly, previously published human motor cortex datasets (Bakken et al., 2021) were projected onto our datasets and heatmap correlation matrices were produced to resolve the last remaining unspecific cluster identities and to confirm our cluster assignments. For the quantitation of fraction nuclei in NC vs HD cell-type specific clusters we utilized full donor datasets for which NeuN negative, NeuN positive / SATB2 negative, NeuN positive / SATB2 medium, and NeuN positive / SATB2 high nuclei could be harvested. Abundance of nuclei in clusters contained within the NeuN positive / SATB2 negative, NeuN positive / SATB2 medium gates were compared between NC and HD samples separately from those clusters contained within the NeuN positive / SATB2 high gates. Percentage of nuclei in each cluster over all nuclei sampled within the sort-gate was calculated and comparisons between HDs and NCs were performed. Comparisons of quantitation were performed using Bonferroni multiple comparison corrected T-test and results with an adjusted P value < 0.05 were marked significant.

### Cell-Type Specific *HTT*-CAG Repeat Expansion Assay

The protocol for this assay was adapted from Ciosi et al. (Ciosi et al., 2019) (Ciosi et al., 2021). Nuclei were isolated following the protocol procedures described in the previous section. DNA was purified using AllPrep DNA/RNA FFPE Kit (Qiagen, #80234) and concentrated in a vacuum concentrator if required and stored at -20°C until further processing. Before initiation of procedures to perform the cell-type specific *HTT*-CAG repeat expansion assay, DNA sample concentrations were measured with Qubit (ThermoFisher #Q32851) and 2-10 ng of genomic DNA was used as template in PCR. Up to 10 ng of gDNA was amplified in a 20 μl volume using NEBNext® High-Fidelity 2X PCR Master Mix (New England Biolabs, #M0541S) supplemented with 5% dimethyl sulfoxide and barcoded primers specific to *HTT* Exon 1 (0.5 μM each). Samples were added to the primer dilutions and a mix containing DMSO and Q5^®^High-Fidelity DNA Polymerase was added. Samples were PCR amplified using the following settings: (96°C – 1min, 30 x (96°C - 30sec, 61°C – 45sec, 72°C – 3 min), 72°C – 10min). After the PCR reaction, 10μl of ddH2O was added to each sample in 20μl and small pools were prepared with the goal for each sample to be represented at similar input concentrations within the pool. Purification was performed by adding 0.55 X AMPure beads. Samples were incubated for 10 minutes, transferred to a magnet, and incubated for 5 minutes. Supernatant was removed and beads were washed twice with 80% EtOH. Beads were dried for 5 minutes and resuspended in 32μl RNase-nuclease-free H2O. Samples were run on Tapestation to check for primer-dimers and the purification process was repeated if necessary. Pool concentrations were assessed by qPCR, using Collibri™ Library Quantification Kit (ThermoFisher # A38524100). Final pooling was based on the concentrations as measured by the Collibri quantification method and samples were sequenced on an Illumina MiSeq Sequencer using a MiSeq Reagent Nano Kit with 30% PhiX spike-in control and 400 nt forward read with output of both indexes.

### Cell-Type Specific *HTT*-CAG Repeat Expansion Data Analyses

Data derived from the Cell-Type Specific *HTT*-CAG Repeat Expansion assay was processed following the steps as outlined by Ciosi et al. (Ciosi et al., 2021). In brief, all HD donors were genotyped for the exact structure ((CAG)n(CAACAG)n(CCGCCA)n(CCG)7(CCT)n or ((CAG)n(CAACAG)n(CCGCCA)n(CCG)10(CCT)n). Sequences derived from the Cell-Type Specific *HTT*-CAG Repeat Expansion Assay were aligned to the appropriate custom-made reference sequences according to the donor’s genotype (CCG)7 or (CCG)10). The number of reads aligned to the reference sequences carrying between 1 and 120 CAG repeats was counted and R code was written to generate result outputs in the form of text files and figures. Mean, standard deviation, minimum, maximum, and range of CAG expansion beyond the inherited mutant allele length were calculated for all cell-type samples from motor cortex.

### *HTT* CAG tract sizing

Demultiplexed sequencing reads were aligned to a set of *HTT* exon 1 reference sequences that differed by the number of CAG repeat units in the repeat tract using the Burrows-Wheeler Aligner (BWA MEM default settings except: -O 6,6 -E 4,4). The number of reads uniquely mapped to each of the reference sequences in the set was considered to reflect the distribution of CAG tract lengths in the two *HTT* alleles of the analyzed cell population. *HTT* read mapping data from each donor was inspected manually for determining the nucleotide sequence of the adjacent polyproline tract and the presence/absence of interruptions in CAG tract. If *mHTT* exon 1 structure of the donor was atypical, then sequencing reads were realigned to a set of reference sequences matching that *mHTT* exon 1 structure. The length of CAG repeat tracts reliably mapped was limited to 113 repeat units. The ratio of somatic expansions (RoSE) (Ciosi et al., 2019), frequency of *mHTT* copies expanded by plus 1 to 2, plus 3 to 5, plus 6 to 20, and by more than 20 repeat units (RU), as shown in example formula below, as well as mean somatic length gain and standard deviation were calculated as follows:

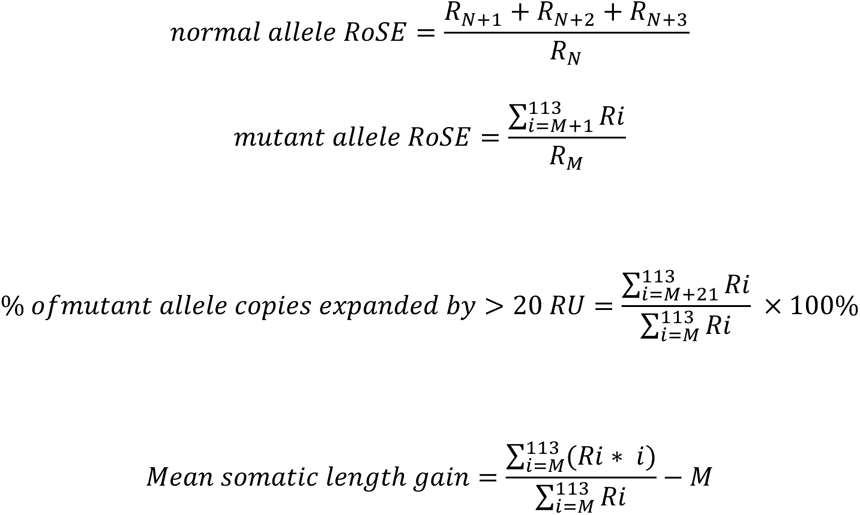

Uninterrupted CAG tract lengths of progenitor/unexpanded *mHTT* allele (M repeat units) and normal *HTT* allele (N repeat units) were defined from the two modes of mapped read length-distribution in CAG-sizing data from non-expanding cell types (for example microglia and astrocytes). R is the number of reads mapped to a reference sequence with the specified CAG tract length. Mean somatic length gain is the average of uninterrupted CAG repeat length in sequencing reads from which the progenitor allele CAG repeat length (M) has been subtracted. It is important to note that the mean somatic length gain does not reflect the increment of size change per mutation event.

